# Antisense transcription from a neighboring gene interferes with the expression of mNeonGreen as a functional *in vivo* fluorescent reporter in the chloroplast of *Chlamydomonas reinhardtii*

**DOI:** 10.1101/2023.11.08.566267

**Authors:** Axel Navarrete, Bernardo Pollak

## Abstract

Although over 30 years have passed since *Chlamydomonas reinhardtii* chloroplast transformation was first achieved, robust genetic engineering of the chloroplast still remains a challenging task. The glass-bead transformation method has enabled simple and accessible chloroplast transformation of the *C. reinhardtii* TN72 strain, allowing generation of marker-free transplastomic strains for low-cost experimentation. However, lack of functional *in vivo* fluorescent reporters limit research and widespread development of chloroplast engineering. Here, we developed a chloroplast codon-optimised mNeonGreen fluorescent reporter, which can be detected *in vivo* through fluorescence microscopy and fluorometry, by context-aware construct engineering, leading up to a ∼6-fold increase in fluorescence. We found evidence for chloroplast post-transcriptional regulation of gene expression derived from formation of antisense pairing of mRNAs due to transcriptional readthrough of the convergent adjacent gene, which was validated through detection of the double-stranded RNA. In addition, engineering approaches were used to modulate transcriptional readthrough, allowing a better understanding of context effects that are relevant for heterologous expression. Finally, we characterised a suite of regulatory parts for transgene expression in a sense-transcriptional context regarding to the selectable marker, achieving up to ∼2-fold increase in mNeonGreen fluorescence levels regarding to the control, with the use of P*rrnS,* 5’*atpA* and 3’*rbcL* endogenous regulatory sequences from the chloroplast of *C. reinhardtii*. This work provides new tools for studying basic aspects of the molecular biology in the chloroplast in *C. reinhardtii*, as well as evidence for fundamental processes of gene regulation that may enable developing rules for more efficient chloroplast engineering.

## INTRODUCTION

Chloroplast transformation was first achieved by biolistics (Boynton et al., 1988; Svab et al. 1990), and is considered the most effective way for delivering DNA into plastids because of its high efficiency. However, high costs of biolistics restricts chloroplast research to well-funded laboratories and limits accessibility. A low-cost chloroplast transformation method using glass beads was developed for *C. reinhardtii* (Kindle et al., 1991) and further optimized by the use of cell wall-deficient strains, which may be used directly without prior digestion of its cell wall (Young & Purton, 2014). This method, along with the use of complementation of an insertional mutant of psbH coding for the photosystem II reaction center protein H in the TN72 strain, enables generation of transplastomic *C. reinhardtii* strains by restoration of photoautotrophy without the need of antibiotics (Wannathong et al., 2016), providing an accessible platform for chloroplast research.

Fluorescent reporters excel in providing detailed spatial and temporal information for the observation of dynamical processes at the cellular and molecular level. Despite substantial progress in the study of the chloroplast’s biology, fluorescent proteins have met limited success as reporters in the chloroplast of *C. reinhardtii*. Previous works have shown the use of several fluorescent proteins including GFP (Franklin et al., 2002; Noor-Mohammadi et al., 2012), mCherry (Kim et al., 2019), the Katushka yellow excitation-red emission protein that showed temporary enhanced fluorescence when chlorophyll was depleted (Suarez et al., 2022), and even the bright Vivid Verde (VFP) (Braun-Galleani et al., 2015) and Venus Q69M (Jackson et al., 2022) fluorescent proteins. However, conflicting evidence has been presented for their poor performance as an *in vivo* reporter, where presence of gene transcripts and protein has been shown but observation of fluorescence through microscopy has been challenging, suggesting that chlorophyll autofluorescence may impede fluorescent reporter visualization (Franklin et al., 2002; Braun-Galleani et al., 2015). In contrast, previous research has shown that proper use of narrow excitation and emission filters can eliminate bleedthrough (Gutiérrez et al. 2022). Furthermore, efforts for reporter fluorescence visualization using nuclear-encoded chloroplast-localized proteins have been successfully undertaken (Karcher et al., 2009; Lauersen et al., 2015; Crozet et al., 2018), suggesting that reporter fluorescence is not an issue in the chloroplast but rather that other events may be affecting chloroplast-encoded fluorescent reporters.

Chloroplasts contain prokaryotic and eukaryotic transcriptional machinery, allowing expression of multiple genes as polycistronic mRNA, as well as possessing transcripts regulated by cis- and trans-splicing (Stern et al., 2010; Barkan, 2011). Transcription in the chloroplast of *C. reinhardtii* is performed by a single plastid-encoded RNA polymerase (PEP), and one nuclear-encoded sigma factor, providing a particular setting for the study of chloroplast transcriptional regulation (Ji et al., 2019). Chloroplast gene expression shows pervasive and genome-wide transcriptional readthrough (Shi et al., 2016), related to inefficient mechanisms for transcriptional termination (Rott et al., 1996), highlighting the importance of RNA degradation in the chloroplast (Schein et al., 2008). New evidence has shown that transgene insertion in the chloroplast can have disruptive effects for native chloroplast genes in the context of insertion (Ghandour et al., 2023). Transgene expression can result in antisense readthrough from adjacent genes, which has been suggested to create untranslatable double-stranded RNA (Rochaix et al., 1989; Prikryl et al., 2011). These reports underscore the need to gain deeper understanding into the biology of chloroplasts, as well as context-aware strategies for more efficient chloroplast engineering.

Available vectors for chloroplast transformation of the TN72 strain, such as the pASapI and derivatives, reestablish the *psbH* functionality and enable expression of a single GOI in an antisense orientation regarding to *psbH* (Braun-Galleani et al., 2015; Wannathong et al., 2016; Young & Purton, 2016; Kim et al., 2020). The architecture of these constructs has been used for expression of recombinant proteins as well as reporter proteins, but observation of fluorescent proteins through microscopy was not reported. Moreover, limited flexibility of these vectors restrict evaluation of transgene orientation effects in heterologous expression.

A system based on Golden-Gate cloning was developed for *C. reinhardtii* chloroplast expression (Bertalan et al., 2015). This system allows modification of regulatory elements, but is tethered to a specific recombination locus and is not interoperable with the Common Syntax for Standard Parts (Patron et al., 2015). Additionally, a modular cloning toolkit for chloroplast biotechnology in higher plants was developed (Occhialini et al., 2019), containing chloroplast regulatory elements, codon-optimized coding sequences for chloroplast expression, and destination vectors suited for chloroplast transformation of *Nicotiana tabacum*, *Zea mays*, and *Solanum tuberosum*. The issue of host-specific requirements for transformation has been approached through the Universal Loop (uLoop) assembly (Pollak et al., 2020). This system has enabled higher flexibility to develop constructs for species-agnostic transformation by decoupling elements required for host transformation, and has been used in bacteria, yeast, plants, diatoms and cell-free systems (Pollak et al., 2020; Taparia et al., 2022., Moosburner et al., 2022; Arce et al., 2021). Moreover, uLoop parts are interoperable with assembly systems that use the Common Syntax, and its versatility could enable the exploring of multiple sequence combinations and their implications for transgene expression in the chloroplast of *C. reinhardtii*.

In this work, we developed vectors for *C. reinhardtii* TN72 chloroplast transformation that enable efficient mNeonGreen (mNG) reporter expression observable through fluorescence microscopy. Different vector designs were tested that included changes in orientation of selectable gene and reporter gene, showing that transcriptional readthrough disrupts expression of the fluorescent reporter when placed in antisense orientation regarding *psbH*. *psbH* evidenced context effects affecting mNG fluorescence levels due to transcriptional readthrough, leading to formation of dsRNA when both genes are in convergent orientation from each other. Confirmation of *mNG* monocistronic transcripts allowed combinatorial approaches of both endogenous and heterologous cis regulatory elements to drive expression of the reporter gene, which were evaluated under different culture conditions to identify functional and efficient combinations for chloroplast transgene expression. Our findings provide new tools for the study of *in vivo* gene expression in the chloroplast of *C. reinhardtii*, and new evidence for post-transcriptional regulation affecting efficient expression of transgenes, highlighting the need for context-aware engineering of the chloroplast.

## RESULTS

A modular vector system for chloroplast transformation of *C. reinhardtii* TN72, based on the pASapI vector (Wannathong et al., 2016), was developed using uLoop assembly (Pollak et al., 2020) (Figure S1, S2a). These vectors consist of four modules including an upstream flanking element (UFE), a fluorescent reporter, the *psbH* transcription unit as a selection marker, and a downstream flanking element (DFE). Two constructs, pLCh1 and pLCh2 vectors demonstrated successful integration and comparable transformation rates to the original pASapI vector (Figure S2b). Additionally, after confirmation of homoplasmicity (Figure S3a), strains exhibited no significant differences in growth rate or transformation efficiency (Figure S4a), rendering them suitable for further experimentation.

### Fluorescent protein expression in chloroplast of *C. reinhardtii*

To evaluate pLCh vectors capability to express a heterologous gene in the chloroplast, pLCh1 and pLCh2 vectors were generated encoding the fluorescent reporter gene *mNG*. Regulatory sequences used to drive expression of *mNG* were cloned from the pASapI vector, and three vectors were generated by modular assembly: a) a pLCh1 vector with the reporter expression driven by the *atpA* promoter (PatpA) and its 5’UTR (Ishikura et al., 1999); a chloroplast codon-optimized *mNG* CDS, and a C-terminal moiety of the *rbcL* coding sequence including its 3’UTR and terminator (pLCh1_mNG::CtRbcL) (Wannathong et al., 2016). b) a pLCh1 vector as in a), but excluding the C-terminal segment of *rbcL* (pLCh1_mNG), as it is suggested to have a minor effect in gene expression (Larrea-Alvarez & Purton, 2020). c) a pLCh2 vector with the same reporter gene as in b), but in the same orientation as the selectable gene (pLCh2_mNG). In addition, a pLCh2_empty vector was used as fluorescence negative control and the four vectors were used for chloroplast transformation of the *C. reinhardtii* TN72 strain. After confirming homoplasmicity (Figure S3b), transplastomic strains were observed by epifluorescence microscopy and their *in vivo* fluorescence intensity was measured by microplate fluorometry.

Epifluorescence microscopy revealed a high signal in the mNG channel of LCh2_mNG strains, which was not seen in other strains (Figure 1a). This signal showed partial co-localization with chloroplast fluorescence visualized in the chlorophyll (Chl) channel (Figure 1a). When visualized through confocal microscopy, the signal detected in the mNG channel of LCh2_mNG strains presented a similar pattern compared to the autofluorescence detected in the Chl channel, but localised in regions with lower autofluorescence inside the chloroplast (Figure 1b).

**Figure 1.**
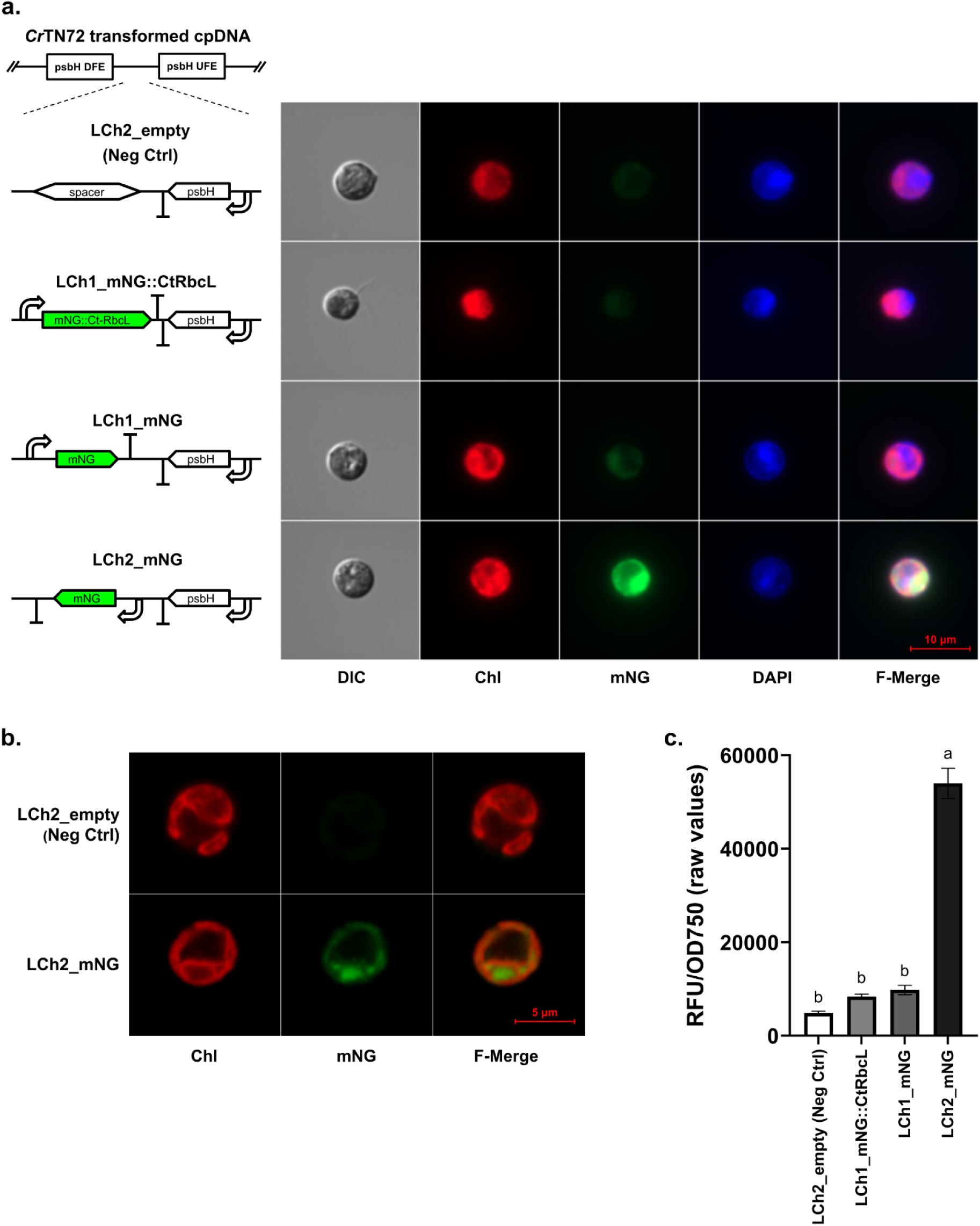
Development of a mNG *in vivo* fluorescent reporter for chloroplast synthetic biology in *C. reinhardtii*. **a)** Fluorescent protein localization in different transplastomic strains determined by epifluorescence microscopy. Insertion unit configurations of the *psbH* locus of each strain are shown in the left. Images show transmitted light (DIC) and fluorescent channels, where chlorophyll autofluorescence (Chl) is represented in red, mNG fluorescence in green, and DAPI staining in blue, respectively. A merged image of all fluorescent channels (F-Merged) was added. Images shown were taken under the same conditions, are unprocessed and were chosen as representatives from 10 different images taken for each strain. **b)** Confocal microscopy showing high-resolution mNG fluorescent protein and its chloroplast localization at higher magnification. Images show different fluorescent channels, where chlorophyll autofluorescence (Chl) is represented in red, mNG fluorescence (mNG) in green, and a merged image of both fluorescent channels (F-Merged). Images shown were taken under the same conditions, are unprocessed and were chosen as representatives from 10 different images taken for each strain. **c)** Relative fluorescence units (RFU) normalized by optical density at 750 nm (OD750) obtained by microplate fluorometry. RFU values were not background subtracted to exhibit that chlorophyll autofluorescence bleeds into the mNG capture range. Error bars represent standard error of mean (SE). Significant differences between groups are depicted with different lower-case letters on top of bars (experimental replicates [n_e_] = 3, biological replicates [n_b_] = 3, technical replicates [n_t_] = 3).

Fluorescence intensity values obtained by *in vivo* fluorometry in transformants obtained with the pLCh1_mNG::CtRbcL and pLCh1_mNG vectors did not present differences that were statistically significant when compared to the negative control, but fluorescence levels of strains obtained by chloroplast transformations using the pLCh2_mNG vector showed statistically significant differences when compared to the negative control and LCh1 strains. LCh2_mNG strains fluorescence values (53962 RFU) were ∼6-fold higher than the other strains (∼9000 RFU) (Figure 1c).

### Effect of the *psbH* terminator in reporter fluorescence

Construct designs used for transgene integration into the chloroplast genome used two transcriptional units adjacent to each other, the reporter and *psbH*. Recent transcriptomic analysis of *C. reinhardtii* chloroplast transcripts revealed that vector designs used for transgene integration in the *psbH* locus use an incomplete *psbH* 3’UTR terminator sequence (TpsbH//) (Gallaher et al., 2018). Due to this, different terminator sequences were tested to replace the incomplete 3’UTR terminator sequence used in the *psbH* selection gene to evaluate transcription conditions of the psbH selectable gene and the reporter gene in strains transformed with pLCh2 vectors (Figure 2a), and were subsequently compared with LCh1_mNG and LCh2_mNG strains. One endogenous and three heterologous 3’UTR + terminator sequences from MoChlo kit (Occhialini et al., 2019) were tested as terminators for *psbH*: the *rbcL* terminator (TrbcL), the *psbA* terminator from *Nicotiana tabacum* (TpsbA(*Nt*)), the tobacco mosaic virus terminator (Ttmv), and the *rrnB* terminator from *E. coli* (TrrnB(*Ec*)), in the pLCh2_H-TrbcL, pLCh2_H-TpsbA(*Nt*), pLCh2_H-Ttmv, pLCh2_H-TrrnB(*Ec*) vectors, respectively. All vectors also contained the same *mNG* transcription unit downstream of *psbH* as pLCh2_mNG. Transformants obtained using the pLCh2_mNG vector were used as positive control to evaluate fluorescence expression.

**Figure 2.**
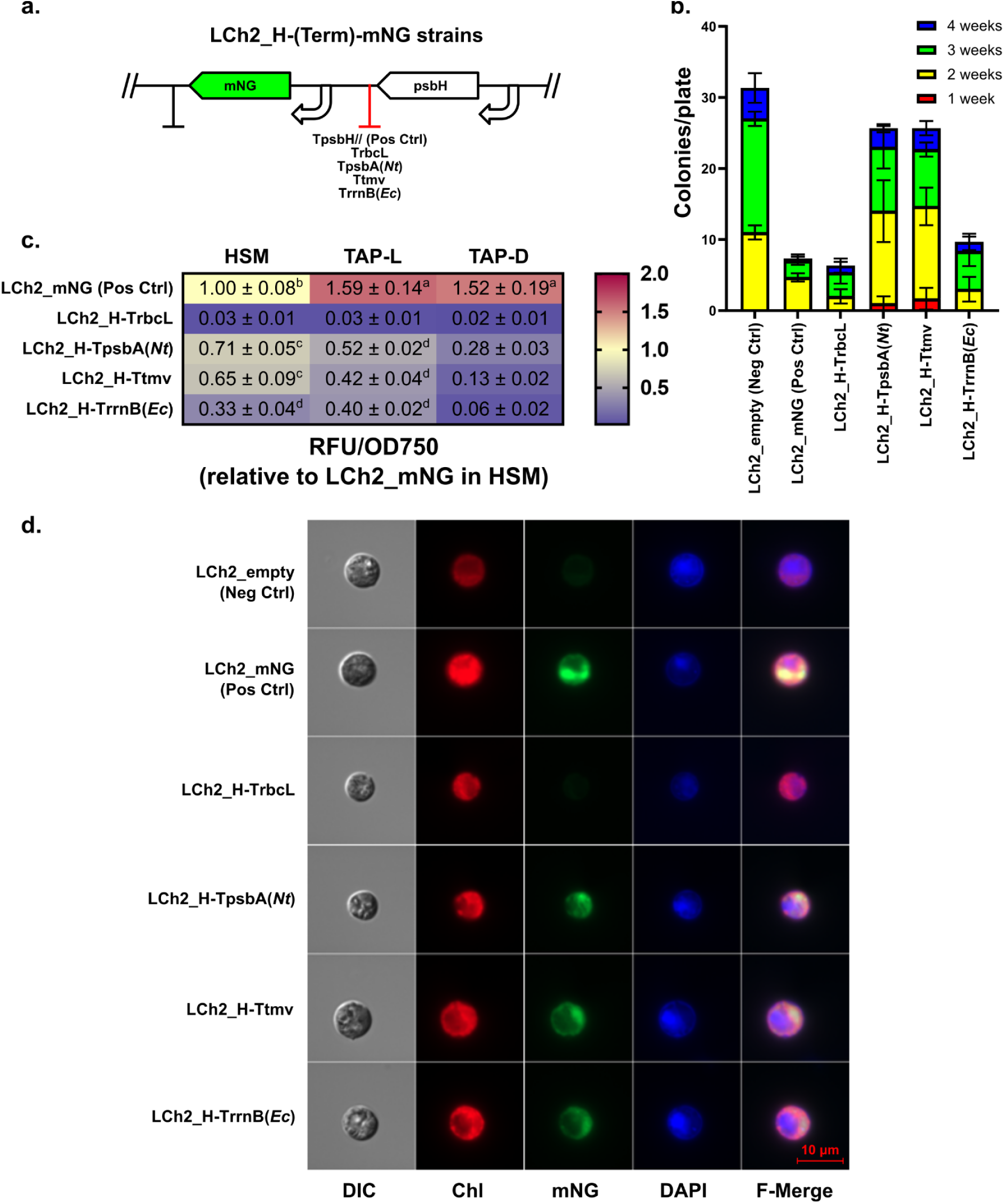
Context effects of selection marker terminator in transformation efficiency and reporter fluorescence levels. **a)** Diagram of the insertion unit of LCh2_H-(Term)-mNG strains in the *psbH* locus. The terminator replaced for modulating transcriptional readthrough of the *psbH* selectable gene is depicted in red. **b)** Number of colonies obtained from glass-bead chloroplast transformation using different LCh2 vectors. Transformant colonies were counted in different weeks after the transformation event. Error bars represent standard error of mean (SE) (experimental replicates [n_e_] = 3, biological replicates [n_b_] = 1, technical replicates [n_t_] = 3). **c)** Heatmap of relative fluorescence units (RFU) normalized by optical density at 750 nm (OD750) obtained by microplate fluorometry of *in vivo* samples of *C. reinhardtii* cultured in photoautotrophic (HSM), mixotrophic (TAP-L) and heterotrophic (TAP-D) conditions. Values are relative to LCh2_mNG strains cultured in HSM conditions and include standard error of mean (SE). Significant differences between groups are depicted with different lower-case letters adjacent to SE, and values without lower-case letters depict that no significant differences were found regarding to negative control strain negative control strain LCh2_empty (experimental replicates [n_e_] = 3, biological replicates [n_b_] = 3, technical replicates [n_t_] = 3). **d)** Fluorescent protein expression in different transplastomic strains determined by epifluorescence microscopy. Images show transmitted light (DIC) and fluorescent channels, where chlorophyll autofluorescence (Chl) is represented in red, mNG fluorescence in green, and DAPI staining in blue, respectively. A merged image of all fluorescent channels (F-Merged) is included. Images shown were taken under the same conditions, are unprocessed and were chosen as representatives from 10 different images taken for each strain.

Transformant colonies were compared among different constructs used, and TpsbA(*Nt*) and Ttmv terminators for *psbH* sequences presented similar transformation efficiency when compared to transformations using the pLCh2_empty vector, and showed an improvement whencompared to the positive control. This did not occur with vectors containing TrbcL and TrrnB(*Ec*) sequences. Notably, transformations with vectors containing TpsbA(*Nt*) and Ttmv sequences presented transformant colonies the first week after transformation (Figure 2b). After confirmation of homoplasmicity of transformed strains (Figure S3c), *in vivo* fluorescence levels of *C. reinhardtii* liquid cultures grown under different conditions were measured by microplate fluorometry, and only strains grown under photoautotrophic conditions were observed through epifluorescence microscopy.

Fluorometry showed that fluorescence levels in positive control strains presented the highest fluorescence intensity among all strains in every growth condition, showing an increase of ∼50% in fluorescence under mixotrophic and heterotrophic conditions (TAP-L and TAP-D, respectively) compared to photoautotrophic conditions (HSM) (Figure 2c). Strains containing terminators different than TpsbH// in the selection gene presented lower fluorescence levels than the control strain in all conditions, with LCh2_H-TpsbA(*Nt*) and LCh2_H-Ttmv strains showing ∼30% less compared to the control, followed by LCh2_H:TrrnB(*Ec*) with a reduction of ∼70% of fluorescence levels; and finally, LCh2_H-TrbcL strains presented no significant differences to negative control strains in any of the culture conditions (Figure 2c). Notably, the positive control maintained high fluorescence levels in heterotrophic conditions, while in the same condition, strains with terminator sequences different than TpsbH// presented the lowest fluorescence intensity among all conditions.

Fluorescence intensity detected through epifluorescence microscopy (Figure 2d) was in agreement with the results obtained by fluorometry. As in LCh2_mNG strains, Almost every strain with *psbH* terminator sequences different than TpsbH// showed fluorescent signals in the mNG channel, with exception of LCh2_H-TrbcL strains, which did not present discernible fluorescence. Fluorescent signals in the mNG channel from LCh2_H-TpsbA(*Nt*), LCh2_H-Ttmv and LCh2_H-TrrnB(*Ec*) strains showed partial co-localization with chloroplast autofluorescence in the Chl channel, as in the positive control strain.

### Analysis of transcript levels, transcriptional readthrough of *psbH* and *mNG* in sense and antisense orientation, and detection of dsRNA in the chloroplast of *C. reinhardtii*

To evaluate underlying causes for the effects observed from the psbH terminator on the reporter gene expression, semiquantitative RT-PCR was performed to find evidence of transcriptional readthrough occurring in *psbH* and *mNG*. Using specific primers for cDNA synthesis through reverse transcription of total RNA from a representative strain of every construct, unterminated transcripts derived from *psbH* and *mNG* transcription were evaluated (Figure 3a). Agarose electrophoresis showed that transcriptional readthrough of *psbH* (R1 and R2) up to the *mNG* coding sequence (*psbH-mNG*) was occurring in all strains except in the negative control (Figure 3b). On the other hand, *mNG* transcripts were detected in both LCh1_mNG and LCh2_mNG strains, but not in the negative control strain (Figure 3b, *mNG* mRNA). In addition, readthrough of *mNG* (*mNG-psbH*) up to the *psbH* CDS (R3v2) was detected in the LCh1_mNG strain, but not in the reverse orientation of transcriptional units (R4) of the LCh2_mNG strain (Figure 3b, *mNG* mRNA). Amplification products for *mNG-psbH* and *psbH-mNG* were verified by Sanger sequencing and sequence alignment confirmed transcriptional readthrough.

**Figure 3.**
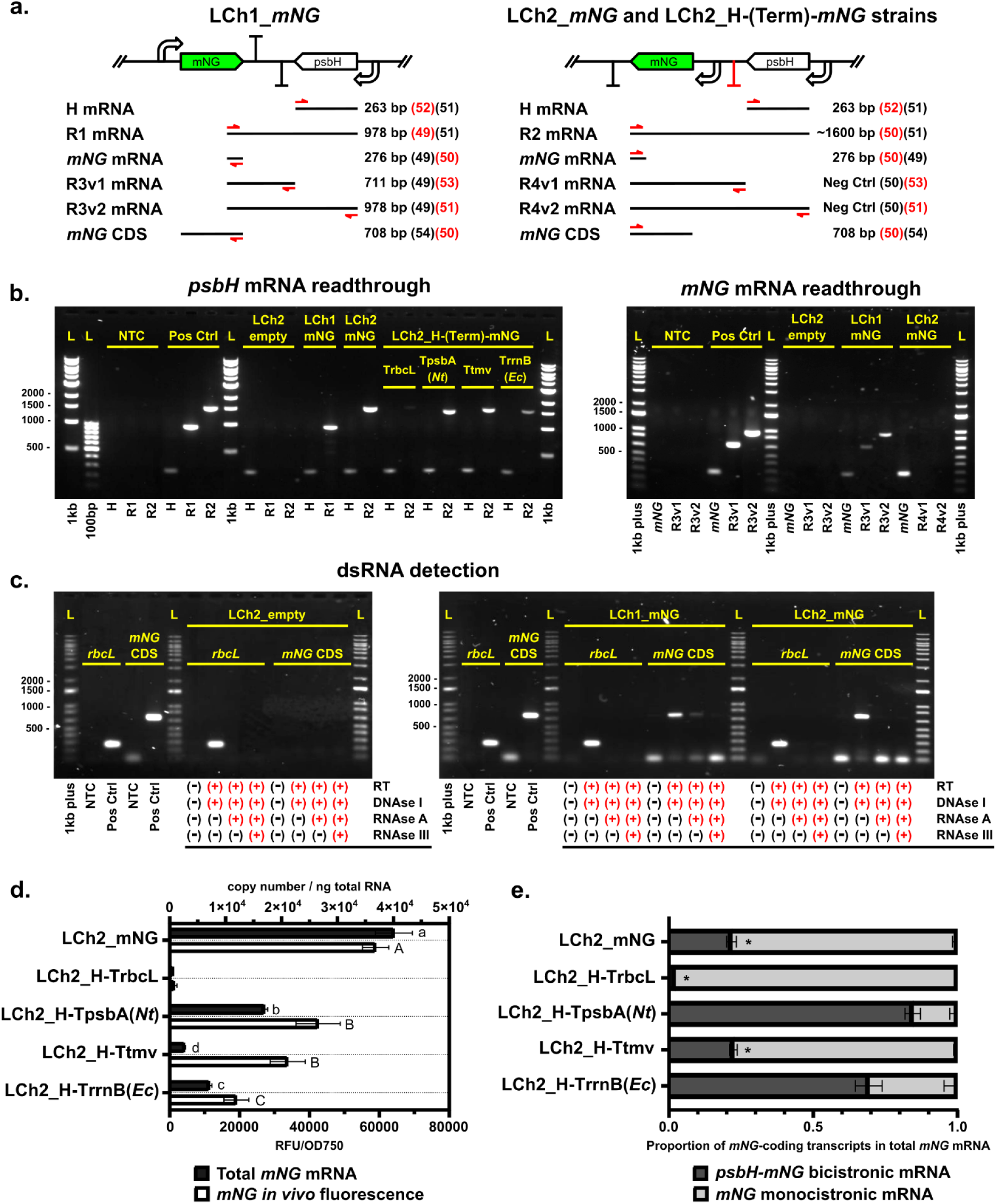
Transcriptional readthrough of *psbH* and *mNG* and detection of dsRNA formed by *psbH-mNG* and *mNG-psbH* transcripts. **a)** Experimental strategy for cDNA synthesis used in *psbH* readthrough, *mNG* readthrough, and dsRNA detection experiments for LCh1 and LCh2 strains. Position of primer alignments for cDNA synthesis by reverse transcription of mRNA are depicted with red directional arrows aligned to the putative transcript. The primers used for reverse transcription are described with its list number in red next to the PCR product size (Supplementary Table 1). For PCR of cDNA, the forward primers used are depicted in black alongside its list number, with the reverse primer being the same used for cDNA synthesis. **b)** Detection of *psbH* mRNA readthrough and *mNG* mRNA readthrough by semiquantitative RT-PCR. cDNAs obtained from total RNA samples of representative transplastomic strains of *C. reinhardtii* were used as PCR templates. Amplicons were separated by agarose gel electrophoresis. Control samples without addition of template are represented as “no template control” (NTC), and positive control templates consisted of gDNA from LCh2_mNG strain (for H and R2 transcripts) and from LCh1_mNG (for *mNG* and R3 transcripts). Ladders used were <u>1</u> <u>kb</u> DNA ladder (NEB), GeneRuler <u>100 bp</u> DNA ladder (Thermo Scientific), and <u>1 kb plus</u> DNA ladder (NEB). **c)** Detection of dsRNA in total RNA samples from representative transplastomic strains of *C. reinhardtii* using ribonuclease treatments and semiquantitative RT-PCR. Amplicons were visualized by agarose gel electrophoresis. Ribonuclease treatments performed prior to each RT-PCR are depicted with (+) at the bottom of the respective lane. PCR products were obtained using cDNA as templates, unless indicated to be negative for reverse transcriptase (RT). Control samples without addition of template are represented as “no template control” (NTC), and positive control samples are shown as “Pos Ctrl”, which templates consist of gDNA from the LCh2_mNG strain. A 264 bp segment of the rbcL gene was used as loading and treatment control. Ladder used was a 1 kb Plus DNA ladder (NEB). **d)** Determination of total *mNG* transcripts through absolute quantification of transcripts by RT-qPCR, and its contrast with *in vivo* fluorescence values obtained for each respective strain. Error bars represent standard error of mean (SE). Significant differences between groups are depicted with different lower-case letters on top of bars for transcript levels, and upper-case letters for *in vivo* fluorescence values (experimental replicates [n_e_] = 3, biological replicates [n_b_] = 1, technical replicates [n_t_] = 3). **e)** Determination of proportion between *mNG* monocistronic transcripts and *psbH-mNG* bicistronic transcripts in relation to total *mNG* transcripts through absolute quantification of transcripts by RT-qPCR. Error bars represent standard error of mean (SE). Asterisks remark values with significant differences between *psbH-mNG* bicistronic transcripts and total *mNG* transcripts in the same sample. Calculation of *mNG* monocistronic transcripts was made by subtraction of *psbH-mNG* bicistronic transcript values from total *mNG* transcript values (experimental replicates [n_e_] = 3, biological replicates [n_b_] = 1, technical replicates [n_t_] = 3).

To determine that formation of dsRNA was occurring due to antisense pairing of *psbH-mNG* and *mNG-psbH* transcripts in the LCh1_mNG strains, detection of dsRNA through semiquantitative RT-PCR was performed using samples treated with RNAse A + 0.6M NaCl for specific degradation of ssRNA, and samples with an additional digestion with RNAse III for specific degradation of dsRNA (Charoonnart et al., 2019). The presence of dsRNA was confirmed by PCR amplification of a 708 bp product present in samples of LCh1_mNG mRNA digested with RNAse A, which could not be detected when the LCh1_mNG mRNA samples included an additional digestion with RNAse III. This did not occur with LCh2_empty and LCh2_mNG strains (Figure 3c).

To evaluate *mNG* expression levels in fluorescent strains, determination of total *mNG* mRNA in the LCh2 strains was performed by absolute quantification of transcripts through RT-qPCR (Figure 3d). LCh2_mNG strains presented the highest level of *mNG* transcripts among all strains (4.01*10^4^ copy number/ng total mRNA), followed by the LCh2_H-TpsbA(*Nt*) strain that presented ∼2-fold lower *mNG* transcript levels (1.69*10^4^ copy number/ng total mRNA) than the LCh2_mNG strain, the LCh2_H-TrrnB(*Ec*) strain that presented ∼4-fold lower *mNG* transcript levels (0.71*10^4^ copy number/ng total mRNA) than the LCh2_mNG strain, and the LCh2_H-Ttmv strain that presented ∼12-fold lower *mNG* transcript levels (0.26*10^4^ copy number/ng total mRNA) than the LCh2_mNG strain. The LCh2_H-TrbcL strain presented *mNG* transcript levels (0.06*10^4^ copy number/ng total mRNA) with no statistical differences from the pLCh2_empty negative control strain.

Due to confirmation of transcriptional readthrough in *psbH* transcripts, determination of *psbH-mNG* bicistronic transcript levels was performed by absolute quantification through RT-qPCR in LCh2 strains. Differences between the amount of *psbH-mNG* bicistronic and total *mNG* transcripts can evidence the presence of *mNG* monocistronic transcripts, since total *mNG* transcripts consider both *psbH-mNG* bicistronic and *mNG* monocistronic transcripts. Absolute quantification of total *mNG* transcripts showed significant differences when compared to *psbH-mNG* bicistronic transcripts in LCh2 strains with TpsbH// (LCh2_mNG), TrbcL and Ttmv 3’UTR + terminator sequences, with copies/ng RNA of total *mNG* transcripts being ∼5-fold higher than *psbH-mNG* bicistronic transcripts in the case of the LCh2_mNG and LCh2_H-Ttmv strains (Figure 3e). This difference in total *mNG* transcripts was ∼100-fold higher than *psbH-mNG* bicistronic transcripts in the LCh2_H-TrbcL strains, but its total *mNG* transcript level was ∼100-fold lower than the LCh2_mNG strains (Figure 3d). Statistical analyses showed there were no significant differences between *psbH-mNG* bicistronic transcripts and total *mNG* transcripts in LCh2_H-TpsbA(*Nt*) and LCh2_H-TrrnB(*Ec*) strains (Figure 3e).

### Determination of fluorescent protein levels in total soluble protein extracts of *C. reinhardtii*

To analyse mNG protein accumulation, total soluble protein (TSP) extracts were obtained and used in native polyacrylamide gel electrophoresis (native-PAGE), and reporter protein presence was evaluated by in-gel fluorescence (Figure 4a), which allowed confirmation of fluorescent protein species without requirement of antibody-dependent techniques (Barahimipour et al., 2015). Gel exposure to blue light showed presence of mNG in protein extracts from LCh1_mNG and LCh2_mNG strains, and also *psbH* terminator variants LCh2_H-TpsbA, LCh2_H-Ttmv and LCh2_H-TrrnB strains; with no visible evidence of mNG in extracts from LCh2_empty and LCh2_H-TrbcL strains. The LCh2_mNG extract presented the highest mNG fluorescence detected, followed by extracts from LCh2_H-TpsbA(*Nt*) and LCh2_H-Ttmv. mNG fluorescence in LCh1_mNG and LCh2_H-TrrnB(*Ec*) extracts was barely detectable (Figure 4a). After fluorescence visualization, the gel was stained with Coomassie Blue, which caused loss of the mNG fluorescence but showed similar amounts of total protein loaded in each well. No migration pattern differences were observed among the samples, nor specific bands for the mNG protein were identified by Coomassie Blue staining (Figure 4a).

**Figure 4.**
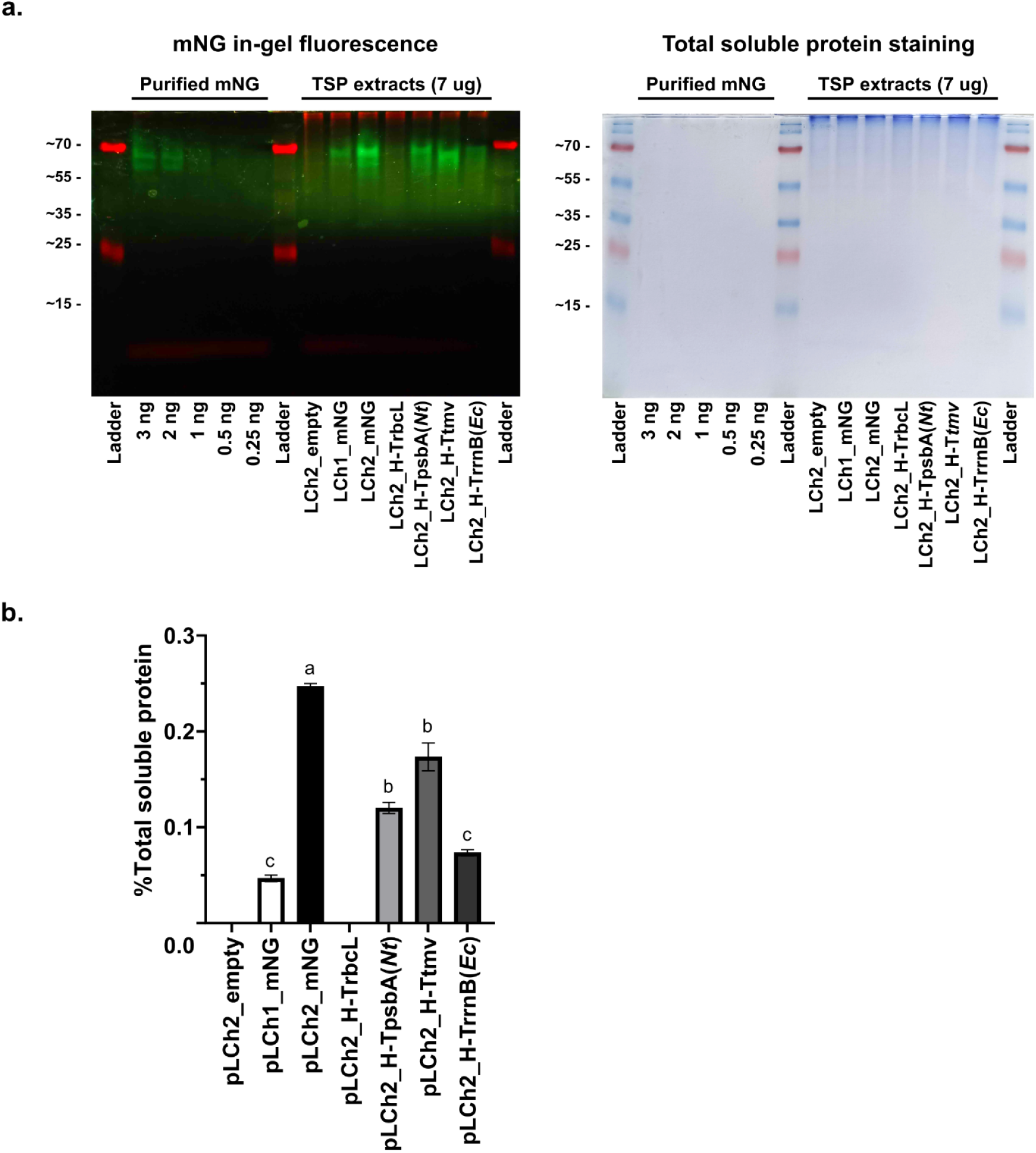
mNG levels in total soluble protein extracts. **a)** Analysis of mNG protein present in total protein extracts of representative transplastomic strains of *C. reinhardtii* by native polyacrylamide gel electrophoresis. Dilutions of purified mNG and 7 µg of total soluble protein extracts obtained from different strains were loaded in each well. Image on the left shows mNG in-gel fluorescence emitted under exposure of blue light and registered by camera using a yellow filter. Image on the right shows the same gel after Coomassie Blue staining to evidence the amount of protein loaded in each well. Ladder used was PageRuler Plus Prestained Protein Ladder (ThermoFisher). **b)** Calculation of mNG protein percentage from total soluble protein extracts of representative transplastomic strains of *C. reinhardtii*. Relative fluorescence units were determined by microplate fluorometry, and fluorescent protein mass was calculated by data interpolation into a standard curve made by dilutions of purified mNG (R^2^=0.98). Error bars represent standard error of mean (SE). Significant differences between groups are depicted with different lower-case letters on top of bars, and values without lower-case letters depict that no significant differences were found regarding to negative control strain LCh2_empty (experimental replicates [n_e_] = 3, biological replicates [n_b_] = 1, technical replicates [n_t_] = 3).

After confirmation of mNG presence in TSP extracts, determination of mNG amounts as percentage in TSP was performed using fluorometry, using purified mNG to generate a standard curve. LCh2_mNG TSP extracts presented the highest levels of mNG accumulation (0.24 %TSP), followed by TSP extracts from LCh2_H-Ttmv and LCh2_H-TpsbA(*Nt*) strains (0.17 and 0.12 %TSP, respectively), and extracts from LCh1_mNG and LCh2_H-TrrnB(*Ec*) presented the lowest levels of mNG accumulation (0.07 and 0.04 %TSP, respectively). mNG amounts in TSP extracts from LCh2_empty and LCh2_H-TrbcL strains were not calculated due to lack of fluorescence in TSP extracts (Figure 4b).

### Characterisation of regulatory elements through reporter fluorescence levels in the chloroplast of *C. reinhardtii*

Due to presence of *mNG* LCh2_mNG and LCh2_H-Ttmv strains, characterisation of promoters, 5’UTR and 3’UTR+terminator combinations from various sources was performed in order to enhance fluorescent reporter expression in the chloroplast. Chloroplast regulatory sequences from heterologous sources, which were obtained from the MoChlo kit (Occhialini et al., 2019), were selected to evaluate their functionality and to avoid unintended homologous recombination to the *C. reinhardtii* chloroplast genome. The pLCh2_H-Ttmv vector design was used for this experiment and named pLCh2t, due that it allows *in vivo* fluorescence, has a higher ratio of mNG protein over transcript, and that transformant colonies can be obtained after the first week of selection. Modifications of the regulatory elements in this experiment consisted in replacing one or two regulatory elements per construct (promoter, or 5’UTR or 3’UTR+terminator) with the mNG transcription unit using the P*atpA,* 5’*atpA* and T*rbcL* regulatory elements as reference (Figure 5a). Selected promoters were: P*atpA*, P*rrnS*, P*psbA*(*Nt*), P*rbcL*(*Nt*), *rpoB*(*Nt*); 5’UTRs: *atpA* 5’UTR, *psbA* 5’UTR, *psbA*(*Nt)* 5’UTR; and 3’UTR+terminators: T*rbcL*, T*rbcL*(*Nt*), T*tymv*, T*rrnB*(*Ec*), and combinations used and construct names are described in (Figure 5b). Generated constructs were used for chloroplast transformation and strains were cultured in photoautotrophic (HSM), mixotrophic (TAP-L) and heterotrophic (TAP-D) conditions and evaluated by fluorometry after confirmation of homoplasmicity (Figure S3d).

**Figure 5.**
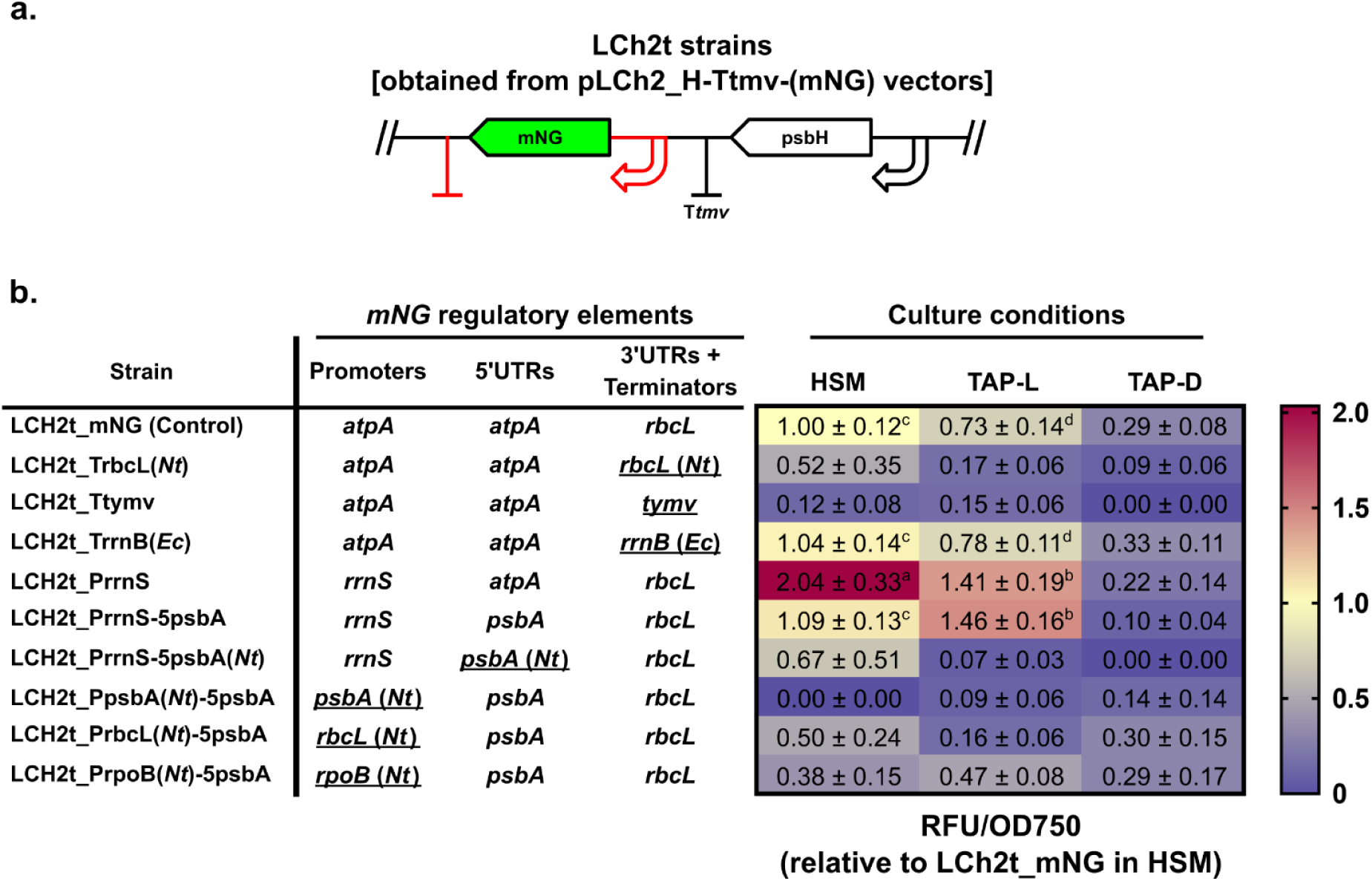
Characterisation of promoters, 5’UTRs and 3’UTR + terminators combinations for reporter gene expression in the LCh2t sense-oriented context in the chloroplast of transplastomic *C. reinhardtii* strains. **a)** Diagram of the insertion unit of LCh2t strains in the psbH locus. Characterisation of regulatory elements affecting expression of the gene of interest are depicted in red. **b)** Heatmap of relative fluorescence units (RFU) normalized by optical density at 750 nm (OD750) obtained by microplate fluorometry of *in vivo* samples of *C. reinhardtii* cultured in photoautotrophic (HSM), mixotrophic (TAP-L) and heterotrophic (TAP-D) conditions. Details of the regulatory elements used in the reporter transcriptional unit for each strain is specified on the left. Heterologous sequences used are shown underlined. Fluorescence values are relative to LCh2t_mNG strains cultured in HSM conditions, and include standard error of mean (SE). Significant differences between groups are depicted with different lower-case letters adjacent to SE, and values without lower-case letters depict that no significant differences were found regarding to negative control strain LCh2_empty (experimental replicates [n_e_] = 3, biological replicates [n_b_] = 3, technical replicates [n_t_] = 3).

Changes to the regulatory sequences of the *mNG* transcription unit showed increased fluorescence levels in LCh2t_TrrnB(Ec), LCh2t_PrrnS, and LCh2t_PrrnS-5psbA strains. Particularly, LCh2t_PrrnS strains showed a ∼2-fold increase in fluorescence levels under phototrophic (HSM) and mixotrophic (TAP-L) conditions compared to the control strain (LCh2t_mNG) in their respective conditions. Furthermore, the LCh2t_PrrnS-5psbA strains displayed comparable fluorescence levels to LCh2t_PrrnS strains in mixotrophic and heterotrophic cultures, but under phototrophic conditions showed ∼2-fold lower fluorescence (Figure 5b).

Both LCh2t_PrrnS and LCh2t_PrrnS-5psbA strains stood out as the only strains incorporating endogenous regulatory sequences, apart from the control. Regarding LCh2t_PrrnS-5psbA(Nt) strain, it resembled the reporter gene design present in LCh2t_PrrnS-5psbA strain but featured the *Nicotiana tabacum* chloroplast version of *psbA* 5’UTR, which showed an important decrease in its *in vivo* fluorescence values, showing no significant differences regarding to negative control strains (LCh2_empty) in all culture conditions (Figure 5b).

Replacing the endogenous T*rbcL* 3’UTR terminator sequence with heterologous T*rbcL* from *N. tabacum* (LCh2t_TrbcL(Nt) strains) resulted in a high decrease of *in vivo* fluorescence in all conditions. A similar result in all culture conditions occurred when endogenous T*rbcL* was replaced by a terminator sequence from *Turnip yellow mosaic virus* (T*tymv*), leading to no significant fluorescence levels compared with negative control. Additionally, LCh2t_TrrnB(Ec) strains demonstrated stable fluorescence levels compared to the control in all culture conditions (Figure 5b).

Transplastomic strains with reporter genes containing *N. tabacum* promoter sequences PpsbA, PrbcL and PrpoB; alongside a replacement of the endogenous *atpA* 5’UTR by endogenous *psbA* 5’UTR (as in LCh2t_PrrnS-5psbA strains), all showed decreased fluorescence levels under all conditions, with no statistically significant differences when compared to the negative control strains (Figure 5b).

## DISCUSSION

This study describes the development of uLoop-based vector designs that allowed the evaluation of different insertion unit configurations in the *C. reinhardtii* TN72 strain. The use of uLoop overcame the limitations of traditional molecular cloning systems by enabling directional changes in transcriptional units, and modifications in regulatory sequences used for expression of reporter and selectable genes. Furthermore, these elements are interoperable with other modular cloning systems that use the Common Syntax, which enabled the use of available chloroplast heterologous regulatory elements for characterisation. The study underscored the potential of the uLoop system in chloroplast research, particularly its capacity to explore molecular mechanisms, study heterologous elements for gene expression, and scrutinize transcriptional and post-transcriptional events in the chloroplast of *C. reinhardtii*.

### Efficient protein production by using pLCh2 vectors for chloroplast transformation of *C. reinhardtii*

mNG fluorescence was successfully detected *in vivo* through fluorimetry and epifluorescence microscopy in transplastomic strains transformed with pLCh2 vectors. Several attempts have been undertaken to use fluorescent protein genes as reporters in the chloroplast of *C. reinhardtii* (Franklin et al., 2002; Braun-Galleani et al., 2015; Kim et al., 2019; Jackson et al., 2022). These studies describe fluorescent protein presence, but lack evidence showing *in vivo* fluorescent cells using imaging methods, which is attributed to quenching of the absorption capacity of fluorescent reporters by chlorophyll autofluorescence (Franklin et al., 2002; Rasala et al., 2013). In contrast, other works show efficient fluorescence from nuclear-encoded chloroplast-localized fluorescent proteins (Karcher et al., 2009; Lauersen et al., 2015; Crozet et al., 2018), suggesting that chloroplast-encoded fluorescent reporters may have low expression or that more sophisticated strategies are required to ensure detection (Noor-Mohammadi et al., 2012; Fields et al., 2019; Gutiérrez et al., 2022). In this work, we generated a *C. reinhardtii* chloroplast codon-optimised version of *mNG* (Shaner et al., 2013), since its excitation and emission is distinct from chlorophyll autofluorescence, and as it is among the brightest monomeric fluorescent proteins reported to date. The use of pLCh1 vectors recapitulates a transformation event with a pASapI vector due to the identity of their regulatory elements composition, and the mNG fluorescence signal of strains obtained using this vector did not present significant differences compared with negative control strains. However, the change in orientation of the reporter in pLCh2 vectors enabled *in vivo* detection of mNG through fluorometry and epifluorescence microscopy, which suggests issues with transcriptional/post-transcriptional events rather than events related to protein biology.

### Transcriptional readthrough of both selectable gene and reporter gene affects reporter expression in the chloroplast of *C. reinhardtii* LCh1 strains

Experiments to determine transcriptional readthrough of *psbH* showed that it is pervasive, being present in both LCh1 and LCh2 strains. Chloroplast transcription is known to enable polycistronic expression due to its prokaryotic origin (Barkan, 2011), and while *psbH* transcriptional readthrough product in LCh1 is likely to create a dsRNA through antisense RNA pairing with *mNG* transcriptional readthrough product encoded in the opposite strand, in LCh2 it would generate a bi-cistronic transcript that includes *psbH* and *mNG*, enhancing *mNG* expression. Formation of antisense pairs between both transcription units result in formation of dsRNA, which may interfere with translation (Hotto et al., 2012). Detection of dsRNA in LCh1_mNG strains but not in LCh2 strains suggests that post-transcriptional events may affect *mNG* transcripts, leading to inefficient reporter expression. A similar situation has been reported in adjacent convergent genes being expressed in *N. tabacum* chloroplasts, where GFP could only be detected by protein immunoblot, probably due to lack of *in vivo* fluorescence signal (Tangphatsornruang et al., 2011). In addition, translation of *psbH* mRNA might not be impeded by antisense pairing, due that *psbH* is the selectable gene for this strain, being required for cell viability under photoautotrophic conditions.

Mechanisms including post-transcriptional regulation are described to occur in chloroplasts (Kishine et al., 2004; Schein et al., 2008; Sharwood et al., 2011). Among these, Ribonuclease J is one of the major mRNA degradation pathways in prokaryotes, and it has been the only ribonuclease identified to have a role in the chloroplast of *C. reinhardtii* (Liponska et al., 2018). It has been hypothesized that after an endonuclease cleavage of dsRNA, probably performed by the same enzyme, exoribonuclease activity of ribonuclease J may allow degradation of dsRNA 5’ ends, which may prevent accumulation of dsRNA derived from sense and antisense transcription (De La Sierra-Gallay et al., 2008). Inefficient transcriptional termination in the chloroplast of *C. reinhardtii* may cause difficulties in translation of transcripts that are convergently located in opposite strands. In prokaryotes, antisense RNA can repress translation by sequestering Shine-Dalgarno elements or other regulatory motifs (Waters & Storz, 2009). In chloroplasts, translation of transcripts can also be impeded when 5’UTR sequences are engaged in double-stranded structures, which can be relieved through binding of translation activation factors (Rochaix et al., 1989; Prikryl et al., 2011).

### Substitution of *psbH* terminator in LCh2 strains modulates its transcriptional readthrough and reporter protein accumulation

Absolute quantification of transcripts by RT-qPCR allowed us to determine that terminators tested in this work could not completely avoid transcriptional readthrough of *psbH*, supporting the idea that readthrough is pervasive in the chloroplast of *C. reinhardtii* (Rott et al., 1996). Despite this, LCh2_mNG, LCh2_TrbcL and LCh2_H-Ttmv strains presented significant differences between *psbH-mNG* bicistronic transcripts and total *mNG* transcript levels, the latter including *mNG* contained in *psbH-mNG* bicistronic and *mNG* monocistronic transcripts. T*psbH//* and T*rbcL* are endogenous 3’UTR termination sequences from *C. reinhardtii* chloroplast, which may contribute to termination specificity for transcriptional units whose promoter sequences enable transcription through nuclear-encoded polymerases (NEP) and plastid encoded polymerases (PEP), but the lack of 633 bp in the 3’ end of the *psbH* 3’UTR terminator could lead to a decrease in *psbH* termination efficiency, resulting in higher rates of *psbH-mNG* bicistronic transcripts compared to T*rbcL* 3’UTR terminator. Despite the highly reduced *psbH-mNG* transcriptional readthrough compared with other strains, LCh2_TrbcL strains exhibited the lowest mRNA and protein accumulation levels, indistinguishable from negative control strains. This might be due to the use of the same sequence for consecutive transcriptional units, as the *mNG* reporter also contained T*rbcL* at its 3’ end, which has been described by previous research (Larrea-Alvarez & Purton, 2020). In the case of T*psbA*(*Nt*) sequence, it has been described to have a single hairpin structure and to act as regulator of mRNA transcription by interacting with 5’UTR, which seems to highly increase mRNA stability (Katz & Danon, 2002). It is also known that chloroplast 3’UTR sequences improve stability of mRNA through protein interaction with RNA secondary structures, but these structures are not required for efficient transcription termination (Stern & Gruissem, 1987; Manavski et al., 2018). The T*tmv* 3’UTR termination sequence consists of a 190 bp sequence that contains domains which form pseudo-knot tRNA-like structures (TLS) (Van Belkum et al., 1985; Osman et al., 2000). These structures may increase mRNA stability by avoiding 3’ exonuclease degradation of *psbH* mRNA (Sharwood et al., 2011), which may explain that *in vivo* fluorescence could still be detected despite the low levels of total mNG transcripts quantified by RT-qPCR. The T*rrnB*(*Ec*) sequence is a strong terminator containing two domains that form hairpins for factor-independent transcription termination (Brosius et al. 1981; Orosz et al., 1991). Despite this, there were no significant differences between *psbH-mNG* bicistronic and total *mNG* transcripts in these strains, which sustain the hypothesis that this kind of secondary structures do not contribute substantially to transcription termination in *C. reinhardtii* chloroplast (Stern & Gruissem, 1987; Manavski et al., 2018).

Differences were found for transcript levels of LCh2_*mNG*, T*psbA*, T*tmv* and T*rrnB* strains compared with their *in vivo* fluorescence levels detected by fluorometry, and mNG amounts measured in TSP extracts. This might be attributed to transcript stability and half-life, but previous studies have shown that there is no strict correlation between mRNA levels and protein levels in the chloroplast of *C. reinhardtii* (Eberhard et al., 2002; Rasala et al., 2011).

### Characterisation of endogenous and heterologous regulatory elements through fluorescent reporter expression in chloroplasts of *C. reinhardtii*

The pLCh2t vector design was selected to test different reporter gene conformations due to its capability to generate *mNG* monocistronic transcripts and its capability for reporter protein expression as observed in this work. Using these constructs, characterisation of regulatory elements for driving the expression of the *mNG* reporter gene showed that only endogenous regulatory elements were able to efficiently express fluorescent reporters, with the exception of the T*rrnB*(*Ec*). The use of the T*rrnB*(*Ec*) sequence exhibited fluorescence levels compared to the control, and had a similar trend as a functional terminator for the psbH selectable gene in this work. This implies that highly divergent prokaryote terminators are functional in chloroplasts of *C. reinhardtii* likely through conserved mechanisms of factor-independent termination (Orosz et al., 1991).

In the case of heterologous promoters, different sequences from *N. tabacum* were selected from promoters activated exclusively by PEP (P*psbA*), by NEP (P*rpoB*) or by both (P*rbcL*). All heterologous promoters diminished *in vivo* fluorescence levels to non detectable levels. Results obtained in this work agree with the results reported by previous research where heterologous promoters are unlikely to drive gene expression in *C. reinhardtii*’s chloroplast (Gimpel & Mayfield, 2013), which remarks that endogenous regulatory sequences are likely to work better than heterologous sequences. Similar results were obtained using an heterologous psbA (*Nt*) 5’UTR instead of its endogenous version, which was in agreement with evidence suggesting that only endogenous 5’UTR from *C. reinhardtii* could support translation of a D1 protein, which is likely to form a secondary structure recognized by endogenous factors (Gimpel & Mayfield, 2013).

Regarding the endogenous promoters and 5’UTR sequences used in this work, replacing the atpA promoter by PrrnS increased in vivo fluorescence reporter levels ∼2-fold when compared with LCh2t_mNG strains, and this improvement was attenuated in phototrophic conditions when the atpA 5’UTR was replaced with the psbA 5’UTR. This is in agreement with results obtained in other work, which was attributed to auto-attenuation of psbA 5’UTR dependent from the psbA product D1 protein (Rasala et al., 2011).

In summary, our research presents empirical evidence for antisense transcription leading to formation of dsRNA in the chloroplast of *Chlamydomonas reinhardtii*. We have demonstrated that the developed mNG fluorescent reporter is a powerful tool for probing the intricacies of chloroplast transcription and gene regulation. This tool is particularly valuable in the context of genetic engineering, where comprehending and controlling these fundamental processes is crucial. Our study not only contributes to the growing body of knowledge regarding chloroplast transcription mechanisms but also highlights the importance of RNA processing and degradation in developing effective strategies for chloroplast engineering. The use of fluorescent reporters in our experiments underscores its utility in molecular biology, particularly for *in vivo* studies of gene expression dynamics. By focusing on the expression of a fluorescent reporter in the psbH locus, we have encountered and addressed significant challenges associated with the complex biology of chloroplast transcription in Chlamydomonas. These findings are instrumental in formulating design principles for gene circuits in synthetic biology and enhancing the feasibility of chloroplast research. We anticipate that these advancements will not only facilitate future research in chloroplast biology but also broaden the scope and capabilities of genetic engineering within this domain.

## METHODS

### Cell strains, culture conditions and maintenance

The *C. reinhardtii* cell wall deficient strain TN72 cw15 ΔpsbH (Young & Purton, 2014) was obtained from Chlamydomonas Resource Center’s collection (https://www.chlamycollection.org), listed as CC-5168. Handling of cultures was performed according to standard protocols (Harris, 1989). Agar stabs were streaked in 2% agar TAP plates (Phytotech Labs, T8224), re-streaked in 1.2% agar (Winkler, 101150) YTAP (TAP + 0.4% w/w yeast extract) plates. Strains were restreaked every 3-4 weeks. Sueoka’s High Salt medium (HSM) (Phytotech Labs, S7668) was used for liquid culture and vacuum-filtered prior to autoclaving. Cultures were grown heterotrophically using > 10 PPFD, and photoautotrophically or mixotrophically using ∼50 PPFD, with 3000K LED lighting (Baer et al., 2016). Growth conditions used were 100 rpm agitation and 25 °C. OD was measured at 750 nm by spectroscopy to calculate growth rates and predict ODs for transformation.

### Plasmid construction

DNA manipulations were carried out following standard protocols (Green & Sambrook, 2021). The pASapI vector (Economou et al., 2014) was obtained from Chlamydomonas Resource Center and oligonucleotides were designed (Supplementary Table 1) to create modular DNA parts compatible with the uLoop assembly system (Pollak et al., 2020) through PCR products from the pASapI vector or duplex oligonucleotides obtained from Macrogen Inc. (Seoul, Korea). PCR reactions were conducted using Phusion High-Fidelity DNA Polymerase (Thermo Fisher Scientific, F530S) according to the manufacturer’s protocol. Cloning of Level 0 (L0) vectors was performed with PCR products and annealed oligonucleotides purified by isopropanol precipitation (Green & Sambrook, 2017), cloned into the pL0R-mRFP1 vector receiver (Figure S5) using Golden-Gate assembly (Engler et al., 2008). Vectors were verified through Sanger sequencing by Macrogen, or using a Flongle flow cell (Nanopore, FLO-FLG114) in a MinION device (Oxford Nanopore Technologies, Oxford, UK) and a Rapid Barcoding Kit (Nanopore, SQK-RBK114) according to manufacturer’s protocol. L2 transformation vectors were constructed according to the uLoop assembly protocol (Pollak et al., 2020) and different designs were used for arrangement of modules (Figure S1b). Vector containing codon-optimised fluorescent reporter with addition of uLoop-compatible L0 parts was synthesized by Twist Bioscience (South San Francisco, CA, USA). Codon-optimisation was designed using the “most frequent” algorithm of the CSO software (Weiner et al., 2020), which is based on Kazusa’s codon usage table for the *C. reinhardtii* chloroplast (Nakamura et al. 2000). To create negative control strains, the AF-spacer1 (Pollak et al., 2019) containing a non-coding sequence was used instead of the reporter gene.

Chloroplast heterologous regulatory sequences were selected from the MoChlo kit (https://www.addgene.org/kits/lenaghan-mochlo) (Occhialini et al., 2019). Terminators used for *psbH* were T*psbA* from *Nicotiana tabacum*, T*tmv* from Tobacco Mosaic Virus, and T*rrnB* from *Escherichia coli*, from the MoChlo kit; and the T*rbcL* sequence from the *C. reinhardtii*’s, which was cloned from the pASapI vector. Also, three promoters (P*psbA Nt*, P*rbcL Nt*, P*rpoB Nt*) one 5’UTR (*psbA Nt* 5’UTR), and three 3’UTR (T*rbcL*, T*tymv*, T*rrnB Ec*) were selected from the MoChlo kit to assemble promoter, 5’UTR and terminator combinations for characterisation of regulatory elements. Endogenous sequences for P*rnnS* and the *psbA* 5’UTR were cloned directly from *C. reinhardtii* TN72 strain using listed primers (Supplementary Table 1)

Due to antibiotic resistance incompatibility between MoChlo and the uLoop assembly system, L1 odd receiver vectors with ampicillin (pCAampOdd) or chloramphenicol (pCAchlorOdd) resistance were generated through Gibson assembly (Gibson et al., 2009) using uLoop specific primers 39, 40, 41, 42 (Supplementary Table 1), to be used as shuttle vectors for cloning into L2 vectors. Depending on the pLCh design chosen for chloroplast transformation, assembled L1 vectors were used as donors to assemble the Level 2 (L2) pLCh vectors using the pHCe1 receiver vector, a pCAe-1 (Pollak et al., 2020) with a modification in the pBR322 *ori* to enable high-copy number as described for the pUC *ori* (Lin-Chao et al., 1992). pHCe1 vectors were generated by Golden-Gate assembly using AarI and specific primers (Supplementary Table 1). L2 vector designs were based in different arrangements of the pASapI vector (Figure S1b), and were used directly for glass-beads chloroplast transformation of *C. reinhardtii*.

### uLoop assembly, bacterial transformation and plasmid purification

Modular assembly was performed according to Loop/uLoop protocol (Pollak et al., 2019; 2020) (https://www.protocols.io/view/loop-and-uloop-assembly-yxnfxme), with minor adaptations. DNA was quantified by fluorimetry using the Quantus™ Fluorometer (Promega, E6150) with the QuantiFluor® dsDNA System (Promega, E2670) according to manufacturer’s specifications. Concentrations of 5 fmol/µL for donor parts and 2.5 fmol/µL for receiver plasmids were used in assembly reactions. Plasmids were purified using Wizard® Plus SV Minipreps DNA Purification System (Promega, A1470) and validated by restriction digest and sequencing. Plasmid purification from 50 mL bacterial cultures was made using PureYield™ Plasmid Midiprep System (Promega, A2492) for *C. reindhartii*’s chloroplast transformations. Plasmid sequences of chloroplast transformation vectors generated for this work are available in a Zenodo repository (https://zenodo.org/records/10080533).

### Chloroplast transformation and verification of homoplasmicity

Chloroplast transformation was performed using the glass beads chloroplast transformation method (Kindle et al., 1991) with modifications (Economou et al., 2014) for DNA integration into the *psbH* locus by homologous recombination (Figure S3). Exceptions to the protocol were that DNA used for transformation was measured by fluorometry and fixed to be ∼10 µg. Liquid cultures were grown in a BJPX-200B refrigerated shaker incubator (Biobase, Jinan, Shandong, China) at 100 RPM at 25 °C, and transformation plates were incubated at 25°C in a JSR refrigerated incubator modified with 3000K warm white LED lights to provide 200 PPFD, avoiding light saturation conditions (Virtanen et al., 2019).

Transformant colonies were restreaked after 2-3 weeks, 1-2 times on selective media before homoplasmicity was determined. A loopful of cells was used from isolated colonies, resuspended in 180 µL of HSM liquid media and 90 µL were transferred to a new 1.5 mL tube, where total genomic DNA was extracted adding 10 µL of Edwards Extraction Buffer (Kasajima et al., 2004). *C. reinhardtii* cells were homogenized by pipetting softly, and 1 µL was directly used for homoplasmicity confirmation through PCR amplification using a combination of three primers (Supplementary Table 1) for a duplex PCR reaction (Figure S4).

### *In vivo* microplate fluorometry

Strains were grown under photoautotrophic, mixotrophic or heterotrophic conditions to late-log phase before evaluation. Fluorescence levels were measured by microplate fluorometry using a Synergy H1 Hybrid multimode reader (BioTek Instruments, Vermont, USA). Triplicates of 200 uL for each strain cell culture were transferred into Nunc 96-well black/clear bottom plates (Thermo Scientific, 165305). mNG detection was performed using Ex: 505/12 nm and Em: 535/12 nm, with gain set at 120. HSM medium was used for photoautotrophic conditions and TAP medium was used for heterotrophic and mixotrophic conditions. Fluorescence values were normalized on the basis of OD_750 nm_ in cell cultures. Transplastomic strains with absence of fluorescent reporter were used for background subtraction.

### Fluorescence microscopy

Liquid cell cultures under different conditions were grown to late-log phase in photoautotrophic conditions for light and epifluorescence microscopy; and under mixotrophic conditions for confocal microscopy. To visualize structures different from the chloroplast, cells were stained with DAPI with the following method: 5 µL of DAPI solution (Invitrogen, D1306; 0.5 mg ml^−1^ dissolved in 250 mM sucrose) were mixed with 15 µL of late-log phase cell culture (1-5 x 10^6^ cells/mL) and incubated for 5 min in the dark before visualization (Karcher et al., 2009).

Epifluorescence microscopy was performed in an Axio Imager.M2 microscope (Zeiss, Jena, Germany) equipped with Colibri 7 LED light for excitation at 475 nm for Chl channel, 511 nm for mNG channel, and 385 nm for DAPI channel, with corresponding filters used for Chl (Ex. 480/30 nm, Em. 600 nm longpass), mNG (Ex. 495/20 nm, Em. 540/30 nm) and DAPI (Ex. 365 nm, Em. 445/50 nm). Transmitted light was used for wide-field DIC channel. Chloroplast sub-localization of mNG was determined by confocal microscopy in a LSM 880 AiryScan microscope (Zeiss, Jena, Germany), using a 633 HeNe laser for excitation of chlorophyll, a multiline argon laser for excitation (at 458/488/514 nm) of fluorescent protein, and a 405 diode laser 30mW for DAPI. Filters were used for Chl channel (Em. 645 nm longpass), mNG channel (Ex. 470/40 nm, Em. 525/50 nm) and DAPI channel (Ex. 365 nm, Em. 445/50 nm).

### Transcript analysis by semiquantitative RT-PCR and absolute quantification RT-qPCR

Liquid cell cultures (10 mL) were grown to late-log phase in photoautotrophic standard conditions for total RNA extractions using the Spectrum Plant Total RNA Kit (Sigma-Aldrich, STRN50) by following manufacturer instructions with the following modifications. In the disruption step, cells were homogenized into the disruption buffer only by pipetting, since vortexing drastically decreases RNA yield. Total RNA extracts were treated with Turbo DNAse (Invitrogen, AM2238) for gDNA depletion following manufacturer’s instructions. For semiquantitative RT-PCR, cDNA was synthesized from total RNA using M-MLV Reverse Transcriptase (Promega, M1701) using listed primers (Supplementary Table 1). For detection of dsRNA by semiquantitative RT-PCR, specific RNAse digestions were performed prior to cDNA synthesis. Total RNA was treated with RNAse A + 0.6M NaCl during 1 h for specific degradation of ssRNA, and other samples included an additional digestion with RNAse III during 1h for specific degradation of dsRNA (Charoonnart et al., 2019). Obtained cDNA samples were used as template for PCR reactions, and PCR products were separated by agarose gel electrophoresis in an 1% w/v agarose gel (Lonza, 50004) using 100V during 30 min. For absolute quantification of transcripts by RT-qPCR, KAPA SYBR FAST One-Step RT-qPCR kit (Roche, KK4650) was used following manufacturer protocols. Amplification of specific transcript segments from *psbH-mNG* bicistronic transcripts and total *mNG* transcripts from total RNA was performed using a CFX96 Touch Real-Time PCR Detection System (Bio-Rad Laboratories, CA, USA) using listed primers (Supplementary Table 1). The amounts of *psbH-mNG* bicistronic transcripts and total *mNG* transcripts were calculated according to standard curves (R^2^ = 0.991 and R^2^ = 0.986, respectively), which were prepared using 10-fold serial dilutions of purified amplicons obtained from PCRs of the pLCh2*-*mNG vector using primers 47 and 48 (complementary to PatpA) for *psbH-mNG* bicistronic transcripts, and primers 49 and 50 (complementary to the end of the *mNG* CDS) for total *mNG* transcripts. Standard curves and sample reactions were performed in the same thermocycling batch, using the purified amplicon at 10^6^ copies/ng as positive control. C_q_ values obtained from control reactions using different primer pairs without RNA template (NTCs) were subtracted from values obtained in all samples respectively; subsequently, values obtained from the negative control strain (LCh2_empty) with each primer pair were subtracted from all samples values respectively. Amounts of *mNG* monocistronic transcripts were calculated by subtracting *psbH-mNG* bicistronic transcripts from total *mNG* transcripts. Transcriptional readthrough PCR products were sequence verified through Sanger sequencing (Macrogen Inc., Korea).

### Extraction of total soluble protein, mNG purification, native polyacrylamide gel electrophoresis and protein quantification

Total soluble protein extracts were prepared using algal cell pellets obtained from 40 mL late-log phase photoautotrophic cultures. Purified mNG was prepared using BL21 bacterial cells carrying an L1 vector containing a transcription unit harboring the T7 promoter with a mNG CDS fused to a 6xHistidine Tag and the T7 terminator. Bacterial cells were induced at OD_600 nm_ 0.8 with 1mM IPTG in a 20 mL culture and pelleted 16 h after induction. Algal and bacterial pellets were resuspended in 500 μl and 2 mL of Breaking buffer (pH 7.4) respectively. Breaking Buffer was prepared with 40 mM NaH_2_PO_4_ (Merck, 106346), 600 mM NaCl (Merck, 106404), 20 mM imidazole (Sigma-Aldrich, I2399), 1 mM PMSF (Sigma-Aldrich, P7626). Resuspended cells were transferred to a cryotube with 300 mg of 0.1 mm zirconia/silica beads (Research Products International, 9833), and lysed using Mini-Beadbeater-16 (BioSpec, 607) for 10 disruption cycles consisting in 30 sec of agitation followed by 30 sec of ice incubation. Lysates were centrifuged at 17000 x g for 20 min at 4°C for removal of cell debris and beads. Then, the supernatants were recovered and stored at 4°C. For bacterial extracts, mNG purification was performed using HisPur™ Ni-NTA Resin (Thermo Scientific, 88221) using batch method specified by manufacturer’s protocol. Protein amounts were quantified using Coomassie (Bradford) Protein Assay Kit (Thermo Fisher Scientific, 23200) following manufacturer’s instructions. Native polyacrylamide gel electrophoresis (native-PAGE) was performed using a standard protocol (Harlow & Lane, 1988). Resolving gel was prepared at 12% v/v acrylamide, stacking gel at 5% v/v acrylamide, and electrophoresis was performed using 8V/cm during 90 min. For in-gel fluorescence assay (Barahimipour et al., 2015), excitation of fluorescent samples in gel was performed with the royal blue LED (440/20 nm) and the yellow longpass filter (500 nm) from the Stereo Microscope Fluorescence system (Nightsea, PA, USA). mNG percentage of total soluble protein (TSP) was quantified by microplate fluorometry using the same settings described for *in vivo* fluorimetry. The amount of mNG present in total soluble protein was calculated by interpolating fluorescence data into a standard curve obtained using serial dilutions of purified mNG quantified by the Bradford Assay (R^2^=0.98). Values were normalised to sample TSP quantified by the Bradford Assay to calculate % of mNG in TSP.

### Statistical Analyses

Statistical analyses were performed with the Prism 8.0.1 statistical package (GraphPad Software Inc., California, USA). Following confirmation of normality and homogeneity of variance, significant differences between different samples with a single culture condition were determined by one-way ANOVA, two-way ANOVA was used for multiple culture conditions; both included Tukey’s multiple comparison post-hoc test, at a 95% confidence interval. For absolute quantification of transcripts, significant differences of transcript levels within the same sample were determined by multiple t-tests, significant differences between different samples were determined by one-way ANOVA, Tukey’s multiple comparison post-hoc test with 95% confidence interval.

## AUTHOR CONTRIBUTIONS

**Axel Navarrete:** Investigation, Methodology, Formal Analysis, Visualization, Writing-Original draft preparation. **Bernardo Pollak:** Conceptualization, Methodology, Supervision, Visualization Writing-Review and Editing, Funding acquisition.

## Notes

https://zenodo.org/records/10080533

### Competing Interest Statement

The authors have declared no competing interest.

### Summary of Updates

Changes to text and figures, including new experimental data and removing previous data of less relevance to the manuscript.

