## Supplementary figures for "Antisense transcription from a neighboring gene interferes with the expression of mNeonGreen as a functional *in vivo* fluorescent reporter in the chloroplast of *Chlamydomonas reinhardtii*"

**SUPPLEMENTAL INFORMATION**

**Figure S1 Legend**

Integration of foreign DNA into the *psbH* locus of *C. reinhardtii* TN72 chloroplast genome by homologous recombination, and strategy for confirmation of homoplasmicity.

**a)** Scheme of DNA integration into the *psbH* locus in the chloroplast genome (cpDNA) of the *C. reinhardtii* TN72 strain. Homologous recombination event of the recombination unit (*psbH* downstream flanking element [DFE], *psbH* upstream flanking element [UFE], *psbH* gene [as selectable gene] and GOI) containing a functional *psbH* gene present in a *Cr*TN72 transformation vector is depicted with dotted lines. Transcriptional units and their orientations are illustrated by arrows. The integration unit (functional *psbH* selectable gene and GOI) replaces the interrupted *psbH* gene (//*psbH*) by the *aadA* gene in the *Cr*TN72 strain.

**b)** Progression of transplastomic strains after a successful transformation event. Untransformed cpDNA copies are depicted in blue. Transformed cpDNA copies are depicted in red. As shown in the image, re-streaking newly obtained transplastomic strains while selective pressure is maintained will promote the loss of untransformed cpDNA copies. Number of re-streakings needed to achieve homoplasmicity will depend on the type of selective pressure, in which commonly one re-streaking is enough to achieve homoplasmicity if re-establishment of *psbH* function is used.

**c)** Diagram of strategy used for detection of homoplasmicity and plasmid DNA integration in transplastomic strains obtained from chloroplast transformation using a *Cr*TN72 transformation vector. A PCR using three different primers in a single reaction is performed. Transcriptional units and their orientations are depicted by arrows. Primer A (black) is designed to align with a sequence located outside from the recombination unit. For detection of untransformed copies of cpDNA, primer B is designed to align to the *aadA* gene CDS (blue); and for detection of transformed copies of cpDNA, primer C is designed to align to a sequence present in the GOI (red).

**d)** Detection of plasmid DNA integration into the cpDNA and confirmation of homoplasmicity of transplastomic *C. reinhardtii* strains through PCR and agarose gel electrophoresis. All PCRs were performed using the same mastermix, which possessed three different primers in which two different PCR products can be expected (duplex-PCR). Ladder used was 1 kb DNA ladder (NEB, N3232) and is represented by an L. Sample from a negative homoplasmic strain (NH), which corresponds to TN72 without transforming, is depicted in blue. Sample from heteroplasmic strain (Het), possessing both transformed and untransformed cpDNA copies, is depicted in black. Sample from positive homoplasmic strain (PH), which possesses all of its cpDNA copies transformed, is depicted in red.

**Figure S2 Legend**

Development of pLCh vectors using uLoop assembly for chloroplast transformation of C. reinhardtii TN72 strain, and determination of their transformation efficiency.

**a)** Design and strategy for development of pLCh vectors based on the pASapI vector. Sequences required for chloroplast transformation contained in the pASapI vector were identified and cloned separately into uLoop L0 vectors. Using the BsaI restriction enzyme, L0 parts were assembled into L1 vector modules containing functional units used in chloroplast transformation. Using the SapI restriction enzyme, four pLCh vectors (pLCh0_empty, pLCh1_empty (Hnat), pLCh1_empty, pLCh2_empty) containing different modules (UFE, Spacer, *psbH* and DFE) in Level 1, positions 1-4 of uLoop vectors, were obtained. The pLCh vectors reproduce the pASapI vector, with relevant variations highlighted in red. Hnat refers to a plasmid version containing a *psbH* transcriptional unit cloned as a single part into an L1 module, replicating the native gene. A spacer consisting of a 200 bp non-coding sequence was used to develop vectors without a GOI. Functional elements in vectors that were modularly assembled are depicted following the Synthetic Biology Open Language (SBOL) standard visual symbols. Colors in vector backbones represent different antibiotic resistances.

**b)** Transformation efficiency obtained from glass-bead chloroplast transformation method using different pLCh vectors, with the use of pASapI vector as control. Transformant colonies were counted the 3rd week after the transformation event. Error bars represent standard error of mean (SE). Significant differences between groups are depicted with different lower-case letters adjacent to SE (experimental replicates [$n_e$] = 3, biological replicates [$n_b$] = 1, technical replicates [$n_t$] = 3).

**Figure S3 Legend**

Confirmation of homoplasmicity and plasmid DNA integration into the chloroplast genome of different transplastomic *C. reinhardtii* TN72 strains.

**a)** Detection of homoplasmicity and plasmid DNA integration in transplastomic strains transformed using pASapI vector and different pLCh vectors. Ladder used was 1 kb DNA ladder (NEB, N3232) and is represented by a L. Samples depicted by number are: 1) TN72 without transformation. 2) Three different ASapI strains (obtained through transformation with the pASapI vector). 3) Three different LCh1_empty(Hnat) strains (with the *psbH* transcriptional unit not modularly assembled). 4) Three different LCh1_empty strains. 5) Three different LCh2_empty strains. Expected size of PCR amplicons are shown in yellow, and primers used to obtain the PCR amplicon are specified in parentheses with numbers listed in Supplementary table 1.

**b)** Detection of homoplasmicity and plasmid DNA integration in transplastomic strains transformed using different pLCh vectors that encode *mNG*. Ladder used was 1 kb DNA ladder (NEB, N3232) and is represented by a L. Samples depicted by number are: 1) TN72 without transformation. 2) Three different LCh1_mNG::CtRbcL strains. 3) Three different LCh1_mNG strains. 4) Three different LCh2_mNG strains. Expected size of PCR amplicons are shown in yellow, and primers used to obtain the PCR amplicon are specified in parentheses with numbers listed in Supplementary table 1.

**c)** Detection of homoplasmicity and plasmid DNA integration in transplastomic strains transformed using pLCh1 vector, and pLCh2 vectors where psbH uses different 3'UTR terminator sequences. Ladder used was 1 kb DNA ladder (NEB, N3232) and is represented by a L. Samples depicted by number are: 1) TN72 without transformation. 2) Three different LCh1_mNG strains 3) Three different LCh2_mNG strains. 4) Three different LCh2_H-TrbcL strains. 5) Three different LCh2_H-TpsbA(*Nt*) strains. 6) Three different LCh2_H-Ttmv strains. 7) Three different LCh2_H-TrrnB(*Ec*) strains. Expected size of PCR amplicons are shown in yellow, and primers used to obtain the PCR amplicon are specified in parentheses with numbers listed in Supplementary table 1.

**d)** Detection of homoplasmicity and plasmid DNA integration in transplastomic strains transformed using pLCh2t vectors for characterization of regulatory elements driving *mNG* expression. Ladder used was 1 kb DNA ladder (NEB, N3232) and is represented by a L. Samples depicted by number are: 1) and 6) TN72 without transformation. 2) Three different LCh2t_mNG strains 3) Three different LCh2t_TrbcL(*Nt*) strains. 4) Three different LCh2t_Ttymv strains. 5) Three different LCh2t_TrrnB(*Ec*) strains. 7) Three different LCh2t_PrrnS strains 8) Three different LCh2t_PrrnS-5psbA strains. 9) Three different LCh2t_PrrnS-5psbA(*Nt*) strains. 10) Three different LCh2t_Ppsba(*Nt*)-5psbA strains. 11) Three different LCh2t_PrbcL(*Nt*)-5psbA strains. 12) Three different LCh2t_PrpoB(*Nt*)-5psbA strains. Expected size of PCR amplicons are shown in yellow, and primers used to obtain the PCR amplicon are specified in parentheses with numbers listed in Supplementary table 1.

**Figure S4 Legend**

Evaluation of photosynthetic growth in different transplastomic *C. reinhardtii* strains

**a)** Optical density at 750 nm (OD750) of different *C. reinhardtii* strains obtained by transformation with pASapI and different pLCh vectors. Cells were cultured under photoautotrophic conditions after 4 days of incubation, and were verified to have approximately 0.05 of $OD_{750}$ at the start of incubation. Error bars represent standard error of mean (SE). Significant differences between groups are depicted with different lower-case letters adjacent to SE (experimental replicates [$n_e$] = 3, biological replicates [$n_b$] = 3, technical replicates [$n_t$] = 3)

**b)** Optical density at 750 nm (OD750) of different *C. reinhardtii* strains obtained by transformation with different pLCh2 vectors. Cells were cultured under photoautotrophic conditions after 4 days of incubation, and were verified to have approximately 0.05 of $OD_{750}$ at the start of incubation. Error bars represent standard error of mean (SE). Significant differences between groups are depicted with different lower-case letters adjacent to SE (experimental replicates [$n_e$] = 3, biological replicates [$n_b$] = 3, technical replicates [$n_t$] = 3).

**Figure S5 Legend**

Controls for analysis of RNA for transcriptional status of *psbH* and *mNG*.

**a)** Reactions without a reverse transcription step (-RT) used as controls for detection of *psbH* mRNA readthrough and *mNG* mRNA readthrough by semiquantitative RT-PCR. Total RNA samples of representative transplastomic strains of *C. reinhardtii* were used as PCR templates, which did not include the reverse transcription reaction. Amplicons were separated by agarose gel electrophoresis. Control samples without addition of template are represented as "no template control" (NTC), and positive control templates consisted of gDNA from LCh2_mNG strain (for H and R2 transcripts) and from LCh1_mNG (for *mNG* and R3 transcripts). Ladders used were 1 kb DNA ladder (NEB, N3232), GeneRuler 100 bp DNA ladder (Thermo Scientific, SM0243), and 1 kb plus DNA ladder (NEB, N3200).

**b)** Integrity of total RNA samples from representative transplastomic strains of *C. reinhardtii*, which were treated with different ribonucleases for subsequent use in semiquantitative RT-PCR. Total RNA samples were separated by agarose gel electrophoresis. Ribonuclease treatments performed in each sample are depicted with (+) at the bottom of the respective lane. Ladder used was 1 kb Plus DNA ladder (NEB, N3200).

**Figure S1**

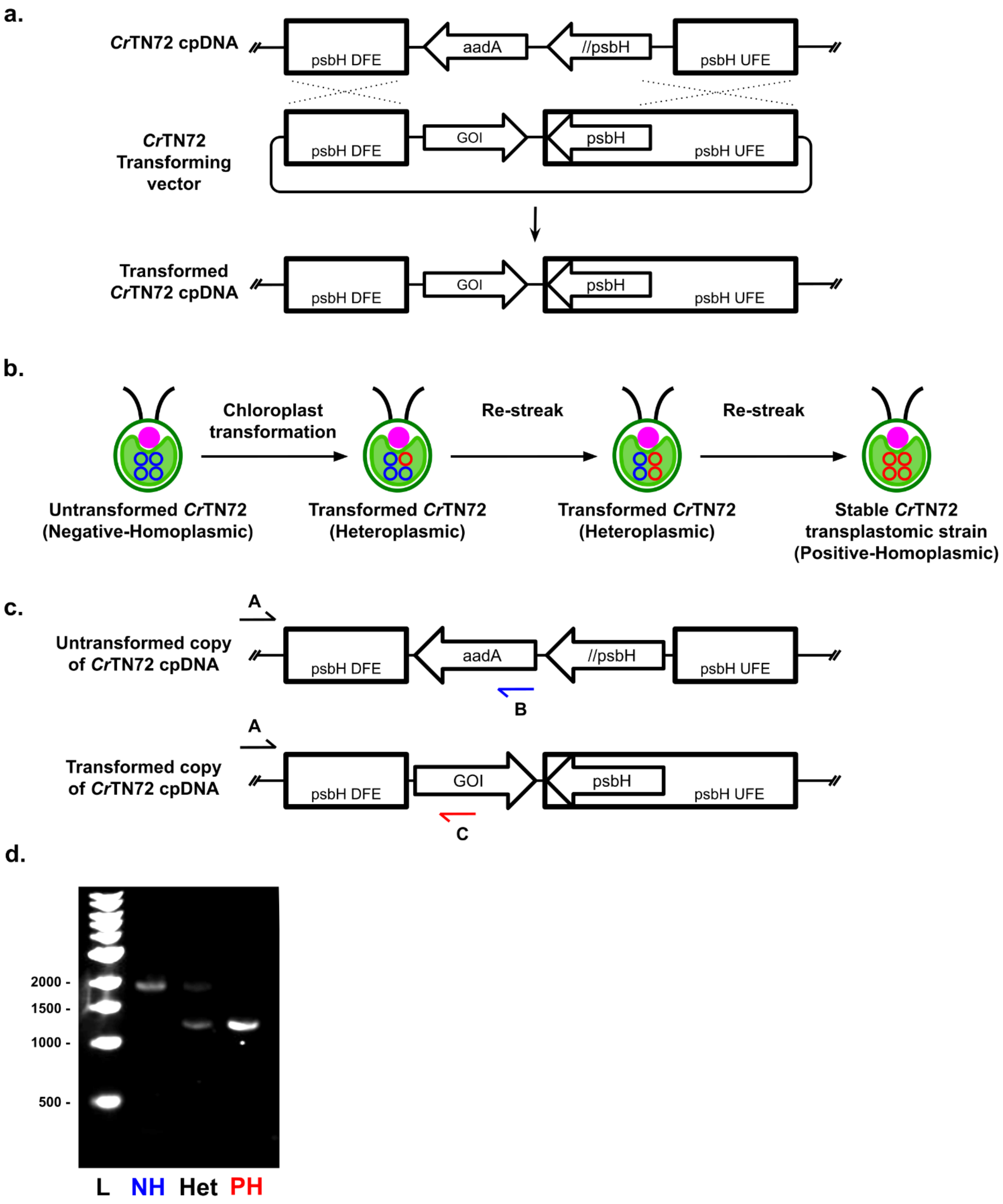

a.
CrTN72 cpDNA
psbH DFE
aadA
//psbH
psbH UFE
CrTN72 Transforming vector
GOI
psbH
Transformed CrTN72 cpDNA
b.
Chloroplast transformation
Re-streak
Re-streak
Untransformed CrTN72 (Negative-Homoplasmic)
Transformed CrTN72 (Heteroplasmic)
Transformed CrTN72 (Heteroplasmic)
Stable CrTN72 transplastomic strain (Positive-Homoplasmic)
c.
A
B
C
Untransformed copy of CrTN72 cpDNA
Transformed copy of CrTN72 cpDNA
d.
2000 -
1500 -
1000 -
500 -
L
NH
Het
PH

#### Figure S2

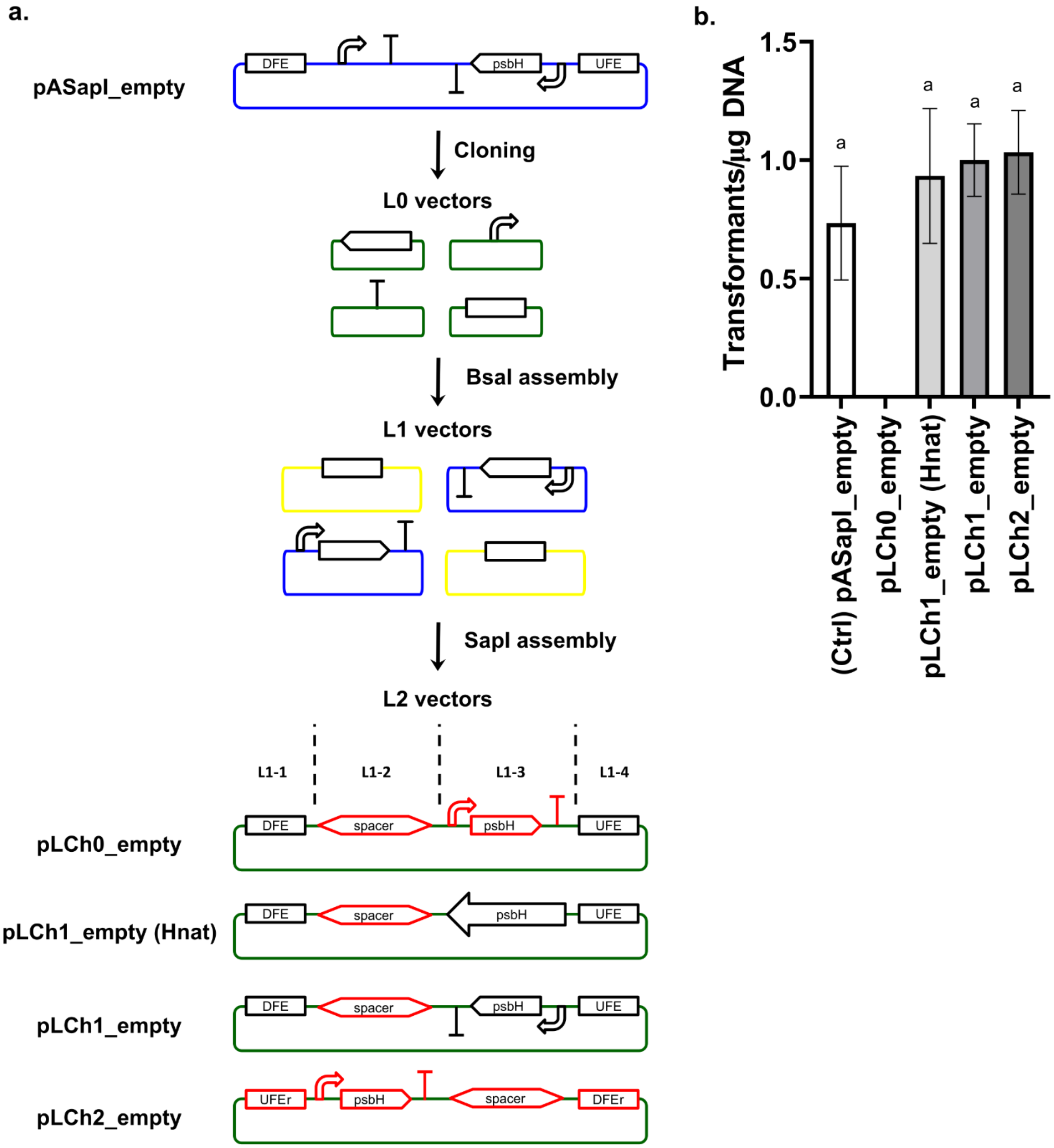

a.
pASapI_empty
DFE
psbH
UFE
Cloning
L0 vectors
BsaI assembly
L1 vectors
SapI assembly
L2 vectors
L1-1
L1-2
L1-3
L1-4
pLCh0_empty
DFE
spacer
psbH
UFE
pLCh1_empty (Hnat)
DFE
spacer
psbH
UFE
pLCh1_empty
DFE
spacer
psbH
UFE
pLCh2_empty
UFEr
psbH
spacer
DFEr
b.
Transformants/µg DNA
1.5
1.0
0.5
0.0
a
a
a
a
(Ctrl) pASapI_empty
pLCh0_empty
pLCh1_empty (Hnat)
pLCh1_empty
pLCh2_empty

#### Figure S3

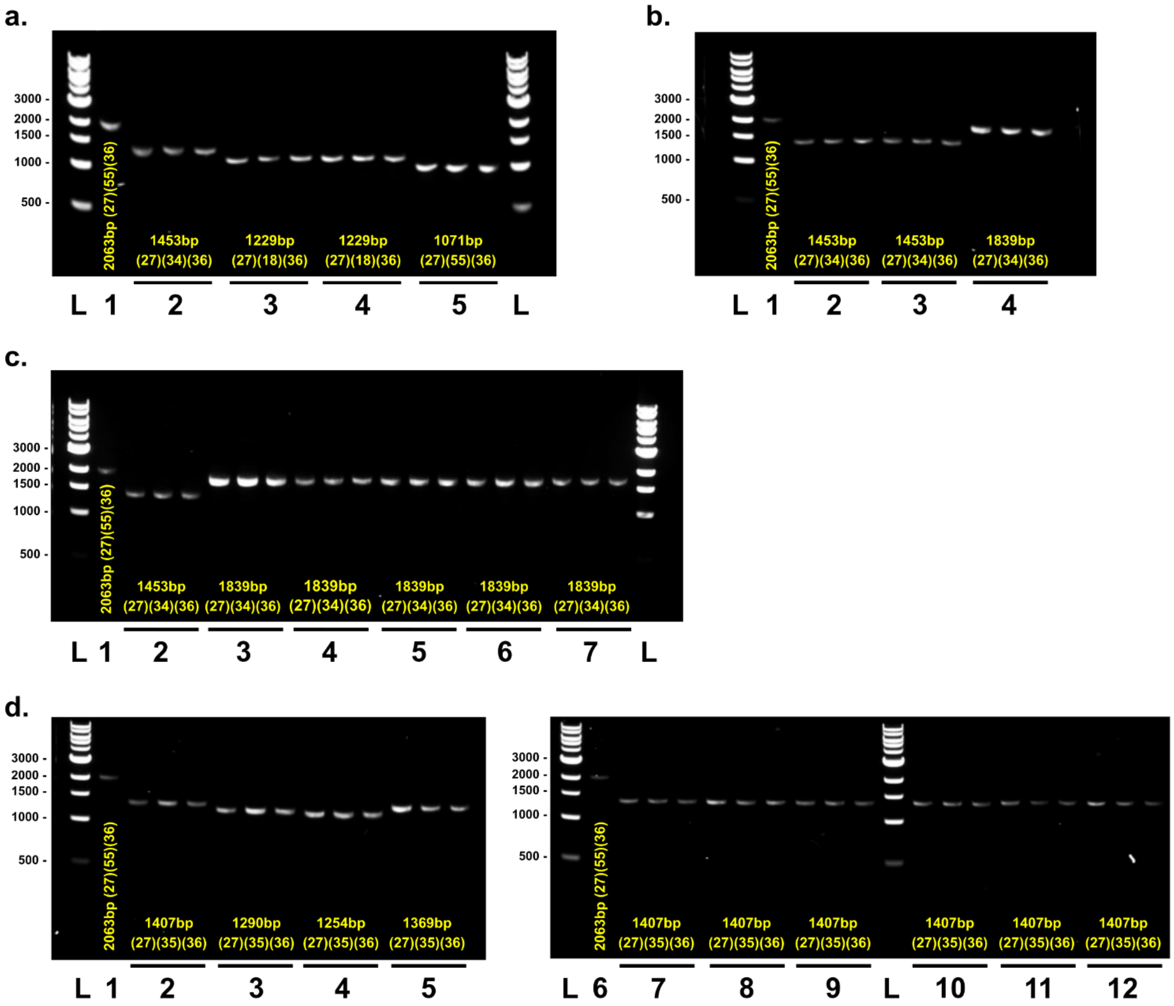
a.
3000 -
2000 -
1500 -
1000 -
500 -
2063bp (27)(55)(36)
1453bp
(27)(34)(36)
1229bp
(27)(18)(36)
1229bp
(27)(18)(36)
1071bp
(27)(55)(36)
L 1 2 3 4 5 L
b.
2063bp (27)(55)(36)
1453bp
(27)(34)(36)
1453bp
(27)(34)(36)
1839bp
(27)(34)(36)
L 1 2 3 4
c.
2063bp (27)(55)(36)
1453bp
(27)(34)(36)
1839bp
(27)(34)(36)
1839bp
(27)(34)(36)
1839bp
(27)(34)(36)
1839bp
(27)(34)(36)
1839bp
(27)(34)(36)
L 1 2 3 4 5 6 7 L
d.
2063bp (27)(55)(36)
1407bp
(27)(35)(36)
1290bp
(27)(35)(36)
1254bp
(27)(35)(36)
1369bp
(27)(35)(36)
L 1 2 3 4 5
2063bp (27)(55)(36)
1407bp
(27)(35)(36)
1407bp
(27)(35)(36)
1407bp
(27)(35)(36)
1407bp
(27)(35)(36)
1407bp
(27)(35)(36)
1407bp
(27)(35)(36)
L 6 7 8 9 L 10 11 12

**Figure S4**

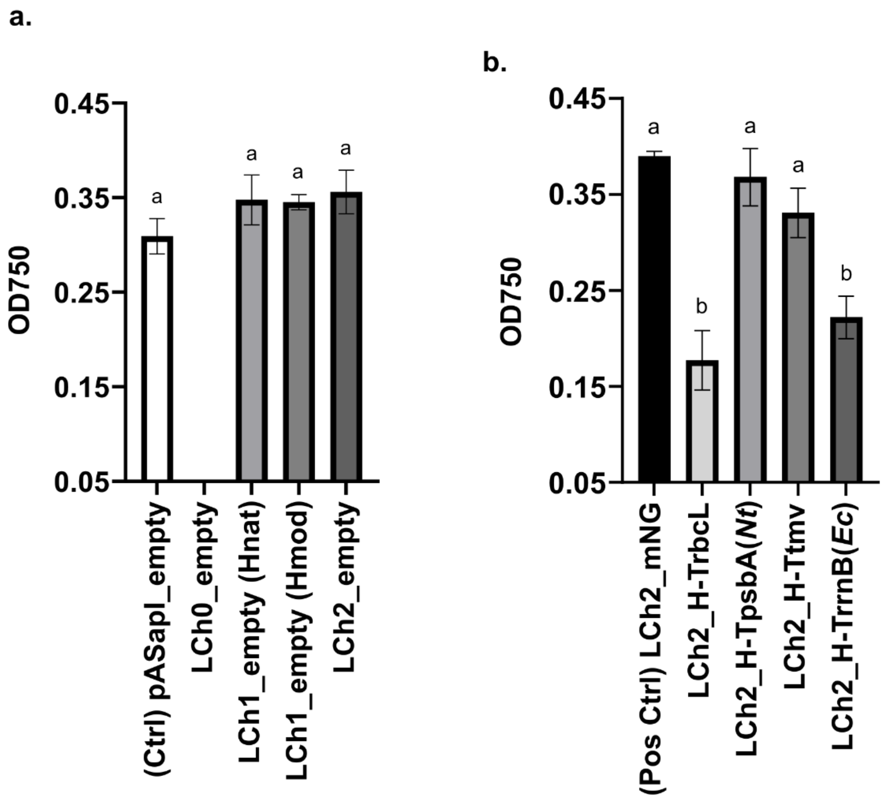
a.
b.
OD750
0.45
0.35
0.25
0.15
0.05
a
a
a
a
(Ctrl) pASapI_empty
LCh0_empty
LCh1_empty (Hnat)
LCh1_empty (Hmod)
LCh2_empty
OD750
0.45
0.35
0.25
0.15
0.05
a
b
a
a
b
(Pos Ctrl) LCh2_mNG
LCh2_H-TrbcL
LCh2_H-TpsbA(*Nt*)
LCh2_H-Ttmv
LCh2_H-TrrnB(*Ec*)

#### Figure S5

a.

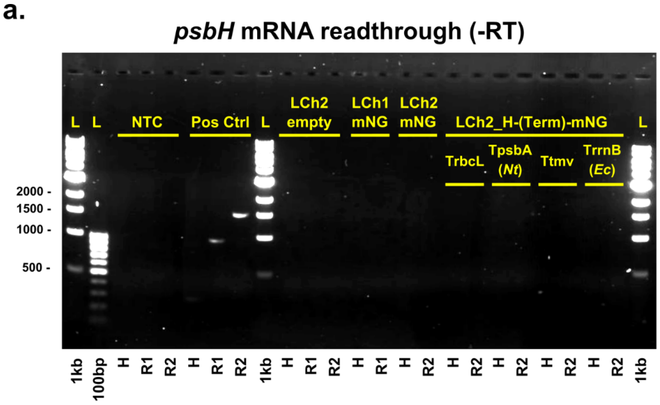
psbH mRNA readthrough (-RT)
L
L
NTC
Pos Ctrl
L
LCh2 empty
LCh1 mNG
LCh2 mNG
LCh2_H-(Term)-mNG
TrbcL
TpsbA (Nt)
Ttmv
TrrnB (Ec)
L
2000 -
1500 -
1000 -
500 -
1kb
100bp
H
R1
R2
H
R1
R2
1kb
H
R1
R2
H
R1
H
R2
H
R2
H
R2
H
R2
H
R2
1kb

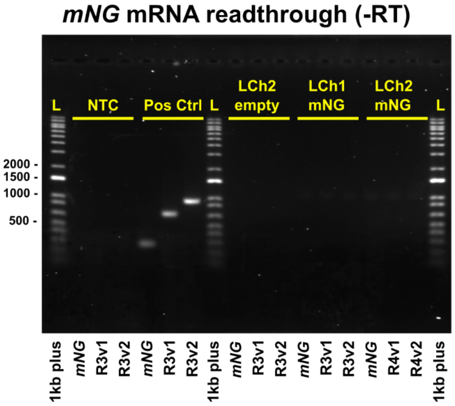
mNG mRNA readthrough (-RT)
L
NTC
Pos Ctrl
L
LCh2 empty
LCh1 mNG
LCh2 mNG
L
2000 -
1500 -
1000 -
500 -
1kb plus
mNG
R3v1
R3v2
mNG
R3v1
R3v2
1kb plus
mNG
R3v1
R3v2
mNG
R3v1
R3v2
mNG
R4v1
R4v2
1kb plus

b.

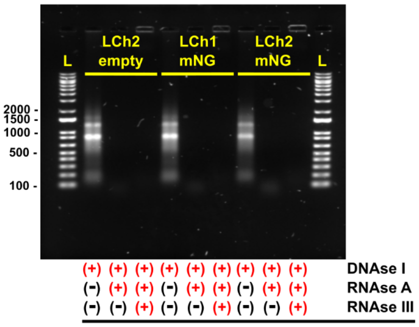
L
LCh2 empty
LCh1 mNG
LCh2 mNG
L
2000 -
1500 -
1000 -
500 -
100 -
(+) (+) (+) (+) (+) (+) (+) (+) (+) DNAse I
(-) (+) (+) (-) (+) (+) (-) (+) (+) RNAse A
(-) (-) (+) (-) (-) (+) (-) (-) (+) RNAse III

### Table S1

#### Table of primers used in this work

| Primer number | Primer name | Sequence | Description |
|---|---|---|---|
| 01 | A_PatpA-F | AAGCTCTTCATCCGGAGacgcgtctccaatatagtagactttattagaggc | For generation of pL0_AB-PatpA from pASapI |
| 02 | B_PatpA-R | TTGCTCTTCTTCGAGTAcatgcaaattaaaaaaaaggtaaatgtatttataaaaagg | |
| 03 | B_5atpA-F | AAGCTCTTCATCCTACTattttaatgcttatgctatctttttatttagtcc | For generation of pL0_BC-5atpA from pASapI |
| 04 | C_5atpA-R | TTGCTCTTCTTCGCATTaaaaagaaaaaataaataaaagattaaaaaagtttatttttaaaa | |
| 05 | dpxDX_TrbcL-F | AAGCTCTTCATCCAGGTtgtactcaagctcgtaacgaaggtcgtgaccttgctcgtgaaggtggcgacgtaattcgttcagcttgtaaatGGaAGAAGAGCAA | For domestication of BsaI restriction site and generation of pL0_DF-TrbcL3'Elem from oligo duplex annealing and pASapI PCR product. |
| 06 | dpxDX_InTrbcL-R | TTGCTCTTCTtCCatttacaagctgaacgaattacgtcgccaccttcacgagcaaggtcacgaccttcgttacgagcttgagtacaACCTGGATGAAGAGCTT | |
| 07 | X_InTrbcL-F | AAGCTCTTCAGGaCTCcagaacttgctgctgc | |
| 08 | F_TrbcL-R | TTGCTCTTCTTCGAGCGtacatccgctttagtatgttactatttcttttattataac | |
| 09 | D_TrbcLv3-F | AAGCTCTTCATCCAGGTtgtactcaagctcgtaacgaaggtcgtg | For generation of pL0_DE-CtailRbcL from an L1 vector containing TrbcL3'Elem. |
| 10 | E_CtailRbcL-R | TTGCTCTTCTTCGAAGCttaaagtttgtcaatagtatcaaattcgaatttaatttc | |
| 11 | E_CTlessTrbcL-F | AAGCTCTTCATCCGCTTtttttatttttcatgatgtttatgtgaatagcataaac | For generation of pL0_DE-CtailRbcL from an L1 vector containing TrbcL3'Elem. Use with primer 8 |
| 12 | A_DFEpsbH-F | AAGCTCTTCATCCGGAGgaatccgcgttttctccgtg | For generation of pL0_AF-DFEpsbH from pASapI. Used for pLCh0 and pLCh1 designs |
| 13 | F_DFEpsbH-R | TTGCTCTTCTTCGAGCGtaatagctcacttttctttaaatttaatttttaatttaaaggtgtaag | |
| 14 | A_PpsbH-F | AAGCTCTTCATCCGGAGgacttgacgactgtgtcccaca | For generation of pL0_AC-PpsbH from pASapI |
| 15 | C_PpsbH-R | TTGCTCTTCTTCGCATTaattgattaaatgaattaagcgttattagcgc | |
| 16 | C_CDSpsbH-F | AAGCTCTTCATCCAATGgcaacaggaacttctaaagctaaacatc | For generation of pL0_CD-CDSpsbH from pASapI |
| 17 | D_CDSpsbH-R | TTGCTCTTCTTCGACCTgaaaccttagctaaagtttcccaactcataga | |
| 18 | E_TpsbH-F | AAGCTCTTCATCCGCTTttttatttaacacaaacataaaatataaaactgtttgttaaggc | For generation of pL0_EF-CDSpsbH from pASapI |
| 19 | F_TpsbH-R | TTGCTCTTCTTCGAGCGttatcttaacggaaggccagtgg | |
| 20 | A_UFEpsbHv2-F | AAGCTCTTCATCCGGAGttttgaactatctaagagatatgttgaa | For domestication of BsaI restriction site and generation of pL0_AF-UFEpsbHv2 from pASapI. Used for pLCh0 and pLCh1 designs |
| 21 | X_InUFEpsbH-R | TTGCTCTTCTtTCttcatggatagacgatagccatttatttacc | |
| 22 | X_InUFEpsbH-F | AAGCTCTTCAGAaACCatatattccagtagcaccgttatga | |
| 23 | F_UFEpsbH-R | TTGCTCTTCTTCGAGCGctgtaagatataaatctaccctgaaagggatg | |

| | | | |
|---|---|---|---|
| 24 | A_psbHRV-F | AAGCTCTTCATCCGGAGttatcttaacggaaggccagtgg | For generation of native oriented psbH TU part pL0_AF-psbHRV from pASapI. Used for pLCh1 designs |
| 25 | F_psbHRV-R | TTGCTCTTCTTCGAGCGgacttgacgactgtgtcccaca | |
| 26 | A_UFEpsbHRV-F | AAGCTCTTCATCCGGAGctgtaagatataaatctaccctgaaagggatg | For generation of pL0_AF-UFEpsbHv2 from pL1-4_UFEpsbHv2. Used for pLCh2 designs |
| 27 | F_UFEpsbHv2RV-R | TTGCTCTTCTTCGAGCGttttgaactatctaagagatatgttgaa | |
| 28 | A_DFEpsbHRV-F | AAGCTCTTCATCCGGAGtaatagctcacttttctttaaatttaatttttaatttaaaggtgtaag | For generation of pL0_AF-DFEpsbH from pL1-1_DFEpsbH. Used for pLCh2 designs |
| 29 | F_DFEpsbHRV-R | TTGCTCTTCTTCGAGCGgaatccgcgttttctccgtg | |
| 30 | A_PrrnS-F | AAGCTCTTCATCCGGAGggcaggcaacaaatttatttattgtcc | For generation of pL0_AB-PrrnS from TN72 gDNA |
| 31 | B_PrrnS-R | TTGCTCTTCTTCGAGTAactctttaaagtttaaattttgtcgggattttaaac | |
| 32 | dpxBC_5psbA-F | AAGCTCTTCATCCTACTgtaccatgcttttaatagaagcttgaatttataaattaaaatattttt acaatattttacggagaaattaaaactttaaaaaaattaacaAATGCGAAGAAGAGCAA | For generation of pL0_BC_5psbA from oligo duplex annealing |
| 33 | dpxBC_5psbA-R | TTGCTCTTCTTCGCATTgttaattttttaaagttttaatttctccgtaaaatattgtaaaaatatt ttaatttataaattcaagcttctattaaaagcatggtacAGTAGGATGAAGAGCTT | |
| 27 | TN72HC_Main-F | GTCATTGCGAAAATACTGGTGC | Used for homoplasmicity checking and GOI insertion into the chloroplast genome |
| 34 | TN72HC_LCh1_Pos-R | aaaaagaaaaaataaataaaagattaaaaaagtttatttttaaaa | Used with TN72HC_Main-F for positive confirmation of homoplasmicity in LCh1 and pASapI strains |
| 35 | TN72HC_LCh2t_Pos-R | GCTGCTGATTGGTGTCGTTC | Used with TN72HC_Main-F for positive confirmation of homoplasmicity in LCh2t strains |
| 36 | TN72HC_Neg-R | ATGgctcgtGAAGCGGTtATCGCC | Used with pASapI_Main-F for confirm absence of GOI integration |
| 37 | L0-F | TGGGCTGCCTGTATCGAGTG | Used for insert verification of L0 parts in Macrogen sequencing |
| 38 | L0-R | GCTGGCCTTTTGCTCACA | |
| 39 | UNS1-F | CATTACTCGCATCCATTCTC | Used for generation of PCR products containing sequence/syntax or backbone from any uLoop vector for Gibson assembly |
| 40 | UNS1-R | GAGACGAGACGAGACAGCCT | |
| 41 | UNSX-F | CCAGGATACATAGATTACCA | |
| 42 | UNSX-R | GGTGGAAGGGCTCGGAGTTG | |
| 43 | HCN1_AarI-U1F | aaaaCACCTGCAACAgagctCATTACTCGCATCCATTCTCAGG | Used for generation of high copy number pHC vectors from any uLoop pCA vector |
| 44 | HCN1_AarI-OriR | ttttCACCTGCAACAtgttcttctagtgtagccgtagttaggc | |
| 45 | HCN2_AarI-OriF | aaaaCACCTGCAACAaacagtatttggtatctgcgctctgc | |

| | | | |
|---|---|---|---|
| 46 | HCN2_AarI-U1R | ttttCACCTGCAACAgctcggcacaaaatcaccactcgatac | |
| 47 | RT-readthrough-F | acgcgtctccaatatagtagact | Used in detection of readthrough transcripts by RT-qPCR |
| 48 | RT-readthrough-R | gagctaaaagagaagaacaatgggt | |
| 49 | RT_mNGCrCp-F | GCTGCTGATTGGTGTCGTTC | Used in detection of total mNeonGreen transcripts by RT-qPCR |
| 50 | RT_mNGCrCp-R | TTTATATAATTCATCCATACCCATAACATCTGT | |
| 51 | RT_psbH-F | gcaacaggaacttctaaagctaaacatc | Used in detection of psbH transcripts |
| 52 | RT_psbH-R | gaaaccttagctaaagtttcccaactcataga | |
| 53 | RT_TpsbH-F | ttttatttaacacaaacataaaatataaaactgtttgttaaggc | Used in detection of readthrough transcripts |
| 54 | RT_mNGCDS-F | atggtttcaaaaggtgaagaagataatatg | Used in detection of mNeonGreen transcripts in RNAse experiments |
| 55 | TN72HC_spc-F | GGGAACAAAGGATTGTGCAGTGCC | Used in confirmation of homoplasmicity in LCh2_empty strains |
| 56 | TN72HC_LCh2_mNG-R | atggtttcaaaaggtgaagaagataatatg | Used in confirmation of homoplasmicity in LCh2_mNG |
